# Hypoxia-inducible factor 1 protects neurons from Sarm1-mediated neurodegeneration

**DOI:** 10.1101/2025.01.17.633664

**Authors:** Paul Meraner, Adel Avetisyan, Kevin Swift, Ya-Chen Cheng, Romina Barria, Marc R. Freeman

## Abstract

The Sarm1 NAD^+^ hydrolase drives neurodegeneration in many contexts, but how Sarm1 activity is regulated remains poorly defined. Using CRISPR/Cas9 screening, we found loss of VHL suppressed Sarm1-mediated cellular degeneration. VHL normally promotes O_2_-dependent constitutive ubiquitination and degradation of hypoxia-inducible factor 1 (HIF-1), but during hypoxia, HIF-1 is stabilized and regulates gene expression. We observed neuroprotection after depletion of VHL or other factors required for HIF-1 degradation, and expression of a non-ubiquitinated HIF-1 variant led to even stronger blockade of axon degeneration in mammals and *Drosophila*. Neuroprotection required HIF-1 DNA binding, prolonged expression, and resulted in broad gene expression changes. Unexpectedly, stabilized HIF-1 prevented the precipitous NAD^+^ loss driven by Sarm1 activation in neurons, despite NAD^+^ hydrolase activity being intrinsic to the Sarm1 TIR domain. Our work argues hypoxia inhibits Sarm1 activity through HIF-1 driven transcriptional changes, rendering neurons less sensitive to Sarm1-mediated neurodegeneration when in a hypoxic state.

**Competing interests:** Marc Freeman is co-founder of Nura Bio, a biotech startup pursuing novel neuroprotective therapies including SARM1 inhibition. The remaining authors declare no competing interests.

## Introduction

Axon degeneration is a hallmark feature of all neurodegenerative diseases but signaling pathways regulating axon degeneration *in vivo* remain poorly defined. After axotomy, axon destruction is driven by the pro-degenerative Sterile alpha and TIR motif containing 1 (Sarm1) molecule. Sarm1 was first identified to be required for Wallerian degeneration in *Drosophila* (dSarm) ^1^, with loss of dSarm blocking distal axon degeneration for weeks after injury. Sarm1/SARM1 is also required for Wallerian degeneration in mouse ^1^ and human neurons ^2^. Extensive work has shown that Sarm1 also promotes axon degeneration in preclinical models of traumatic axonal injury ^3,4^, chemotherapy-induced neuropathy ^5^, diabetes-induced neuropathy ^5^ and ALS ^6–9^. More recently, direct activation of Sarm1 has been demonstrated in response to neurotoxins like Vacor ^10^, further highlighting the broad role for Sarm1 in axonopathies.

How the pro-degenerative function of Sarm1 is activated is best understood in the context of Wallerian degeneration. The Sarm1 TIR domain has intrinsic NAD^+^ hydrolase activity that is constitutively inhibited by its ARM domain ^11,12^. NAD^+^ can bind and stabilize ARM-TIR interactions to keep Sarm1 hydrolase function off, but loss of the axonal survival factor Nmnat2 leads to a drop in axonal NAD^+^ and increase in NMN, the substrate of Nmnat2 ^13^. NMN binds the ARM domain of Sarm1, destabilizes ARM-TIR interactions, and the hydrolase function of Sarm1 is unleashed to drive further NAD^+^ loss. Axon degeneration is thought to result from the catastrophic depletion of axonal NAD^+^ pools by Sarm1. Thus, depletion of Nmnat2, which results in a drop in NAD^+^ and rise in NMN, is considered a key activating event of Sarm1 signaling, and has been proposed to be a widely deployed mechanism for activating Sarm1 in different neurodegenerative settings ^14^. Despite intense study in recent years, major questions remain regarding how Sarm1 activity is regulated, and how it promotes cellular disintegration. Mutations in the ARM domain that potently activate Sarm1 NAD^+^ hydrolase activity and promote degeneration *in vitro* were identified in ALS patients, but why these do not activate neurodegeneration earlier in life remains a mystery ^8^. Loss of the *Drosophila* BTB/Back domain molecule Axed (Axundead) results in complete suppression of dSarm-mediated neurodegeneration, but how Axed mechanistically blocks dSarm pro-degenerative activity remains unclear ^15^, and additional *in vivo* modulators of dSarm/Sarm1 neurodegenerative function await discovery.

To explore how activated Sarm1 drives neurodegeneration, we used an inducible model of Sarm1 activation for genome-wide CRISPR-Cas9 screens in human cell lines. Unexpectedly, we found that ablation of VHL (von Hippel-Lindau disease tumor suppressor) or other substrate receptors or adaptors of Cullin-2 RING E3 ubiquitin ligases (CRL2) suppressed the ability of activated Sarm1 to drive degeneration. A primary target of VHL is the transcription factor hypoxia-inducible factor 1 (HIF-1), which is ubiquitinated with the help of VHL in an O_2_-dependent manner and thereby marked for proteasomal degradation ^16^. Expression of a stabilized, degradation-resistant version of HIF-1 (HIF-1-S) suppressed the ability of activated Sarm1 to drive Wallerian degeneration in mice and flies and resulted in broad gene expression changes in cultured neurons. The neuroprotective effects of HIF-1-S extended to treatment with Vacor (a direct pharmacological activator of Sarm1) or a dimerizable version of the Sarm1 TIR domain, and HIF-1-S expression was sufficient to block the drop in NAD^+^ driven by activated Sarm1. We propose that HIF-1-mediated transcriptional changes potently suppress the ability of Sarm1 to drive neurodegeneration, and this mechanism evolved to mitigate Sarm1 pro-degenerative function in conditions where O_2_ becomes limiting (e.g. stroke or traumatic brain injury), thereby increasing the chances of neuronal survival.

## Results

### Genome-wide CRISPR-Cas9 screens identify Cullin 2-bound substrate receptors as modulators of Sarm1-mediated neurodegeneration

To identify new molecules required for Sarm1 signaling, we performed genome-wide CRISPR-Cas9 sgRNA survival screens in human cell lines. Cells were transduced with doxycycline-inducible SARM1ΔARM **(Supplementary Fig. S1)**, a gain-of-function variant of SARM1 that lacks the autoinhibitory ARM domain and has constitutive NAD^+^ hydrolase activity ^17^. SARM1ΔARM causes rapid neurodegeneration when expressed in primary neurons and leads to cell death in susceptible cell lines. For initial screening the human cell lines HeLa (cervical carcinoma cells) and HEK293T (human embryonic kidney cells) were chosen because of their excellent sensitivity to SARM1ΔARM. To avoid enrichment of false-positive sgRNAs in the screening process – through attenuating mutations introduced into SARM1ΔARM by relatively frequent lentiviral replication errors – we found it essential to first establish clonal lines expressing doxycycline-inducible SARM1ΔARM, verify its chromosomally integrated sequence, and show that SARM1ΔARM induction with doxycycline killed 100% of cells (see methods). Our multicistronic lentiviral expression cassette for SARM1ΔARM **(Supplementary Fig. S1**; see methods) included a EGFP reporter. This feature allowed us to select screen survivors with high levels of EGFP and therefore high levels of SARM1ΔARM, which we reasoned more likely represent SARM1ΔARM-resistant cells that should carry relevant sgRNAs (as opposed to EGFP negative cells that survived by mere suppression of SARM1ΔARM).

Cas9 and sgRNAs were delivered to cell lines in separate transduction steps. In the final libraries, each cell was equipped with three elements: doxycycline-inducible SARM1ΔARM, Cas9, and one sgRNA. Each sgRNA was represented by ∼1000 seed clones, each gene was targeted by 4 orthogonal sgRNAs (a feature of the “Brunello” human sgRNA library ^18^), and the sum of all guides covered the entire genome. Our screens were designed as survival screens, where cells die by default after SARM1ΔARM induction, but are rescued from cell death if harboring a sgRNA directed at a gene that is required for Sarm1 pro-degenerative signaling. Screening results were obtained by analyzing the chromosomally integrated sgRNA repertoire of bulk survivors (**Fig. 1a, left panels)**, and of survivor populations sorted for medium or high levels of EGFP, the reporter of SARM1ΔARM (**Fig. 1a, mid and right panel)**. In our analysis of the screen results, we prioritized genes targeted by several of the 4 orthogonal sgRNAs, and gene families or pathways that were highlighted by high counts for multiple member genes.

**Fig. 1.**
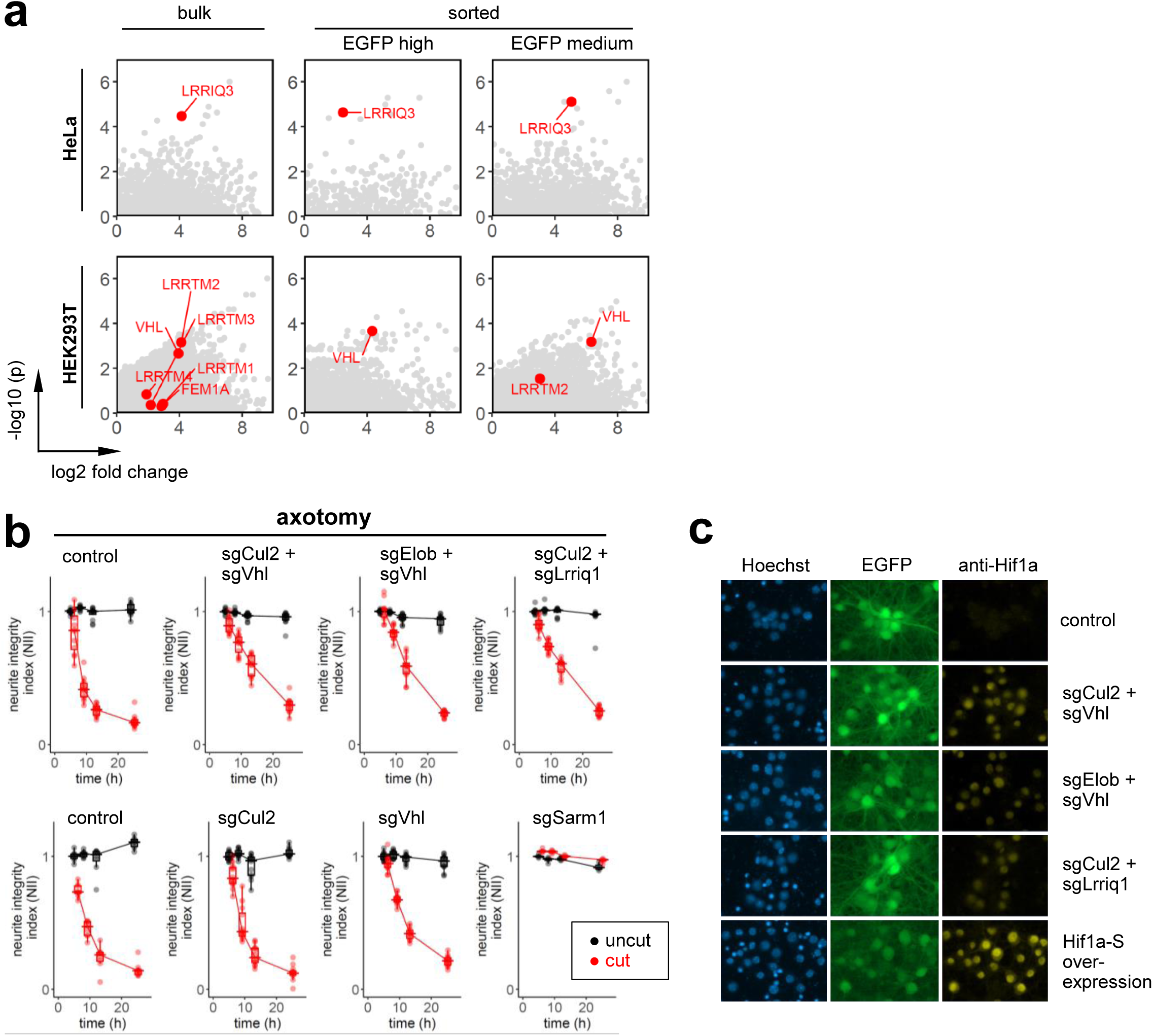
Genome-wide CRISPR-Cas9 screens identify Cullin-2-bound substrate receptors as modulators of Sarm1-mediated neurodegeneration. **a** Volcano plots of sgRNA subsets in cells surviving genome-wide CRISPR-Cas9 sgRNA screens. Cullin-2 substrate receptors of interest highlighted in red. Guides for VHL and LRRIQ3 are also enriched in cell populations with high or medium levels of EGFP – the reporter for SARM1ΔARM – which confirms their status as true positive hits. **b** Axotomy-induced neurodegeneration in mouse Cas9^+/+^ DRG neurons after Cas9-mediated ablation of Cullin-2 and associated adaptors (Elob) and receptors (Vhl, Lrriq1). Axotomy was performed on day 12 after plating, and neurodegeneration was scored using the neurite integrity index (NII, see methods). sgCul2 denotes guides targeting gene *Cul2*; similar notation was used for genes *Vhl*, *Elob*, *Lrriq1*, and *Sarm1*. sgSarm1 was included as a control for Cas9 system performance. **c** Cultured Cas9^+/+^ mouse cortical neurons were stained for HIF-1 alpha chain (Hif1a) after transduction with guide pairs targeting Cullin-2 (sgCul2) and associated receptors Vhl (sgVhl) or Lrriq1 (sgLrriq1), or guides targeting Vhl and canonical adaptor Elob (sgElob). Also shown (bottom) are cells overexpressing a HIF-1 alpha chain variant (Hif1a-S) that cannot be ubiquitinated and degraded. Blue, Hoechst dye staining nuclei; green, EGFP reporter highlighting cell morphology; yellow, Hif1a.

We first focused on the genes VHL (von Hippel-Lindau tumor suppressor) and LRRIQ3 (Leucine-rich repeat and IQ domain-containing protein 3), which showed enrichment with robust scores in the HEK293T or HeLa screen, respectively **(Fig. 1a)**. Both VHL and LRRIQ3 were also found to be enriched in sorted screen survivor populations with high or medium EGFP (acting as SARM1ΔARM reporter) and therefore classified as positive hits **(Fig. 1a, center and right panels)**. Both genes encode known substrate receptors for Cullin-2 RING E3 ubiquitin ligase (CRL2) complexes ^19^, and VHL targets hypoxia-inducible factor 1 (HIF-1) for degradation ^16^ (see **Supplementary Fig. S2a** and **S2b** for CRL2 schematic and components). Under normoxic conditions, the HIF-1 alpha subunit (human gene *HIF1A*) is constitutively hydroxylated by prolyl hydroxylases (human genes *EGLN1, EGLN2, and EGLN3*) at prolines P402 and P564 (P402 and P577 in mouse), using available O_2_. Through hydroxylated prolines, Hif1a is bound by VHL and recruited to CRL2 complexes, ubiquitinated, and proteasomally degraded. In hypoxia, Hif1a hydroxylation is inefficient (due to low O_2_), and the HIF-1 heterodimer is stabilized and accumulates in the nucleus where it acts as a transcription factor, controlling the expression of hundreds of genes ^24,25^. Enrichment of VHL and LRRIQ3 was intriguing, given that prior studies had shown dSarm-mediated neurodegeneration to be blocked by loss of the *Drosophila* BTB/Back domain protein Axundead ^15^, which is a potential bridging molecule for Cullin-3 and its substrates ^20^.

To determine whether VHL and other CRL2-related genes were relevant to SARM1-mediated neurodegeneration, we tested the effects of Cas9-mediated ablation of these genes in primary mouse dorsal root ganglion (DRG) neurons after axotomy (**Fig. 1b**). We found that targeting Cul2 (Cullin-2) or Vhl alone produced minimal neuroprotection (**Fig. 1b, bottom row**). However, dual targeting of Cul2 and Vhl significantly slowed axon degeneration (**Fig. 1b, top row**). Similar protective effects were observed with combined targeting of Vhl and Elob (a canonical CRL2 adaptor) or Cul2 and CRL2 substrate receptor Lrriq1 (**Fig. 1b, top row**), and other combinations (e.g. Cul2 and adaptor Eloc, or Cul2 and Fem1 family receptors; not shown). These results indicate loss of CRL2 substrate receptors or adaptors can suppress both active SARM1ΔARM NAD^+^ hydrolase and Sarm1-dependent neurodegeneration after axotomy in mammalian neurons.

### HIF-1 antagonizes Sarm1-mediated neurodegeneration

Vhl has been extensively studied as a regulator of HIF-1 protein levels ^16,21,22^, and we speculated that sgRNA targeting of CRL2 components might increase stabilized HIF-1 levels as a neuroprotective mechanism. To test this, we transduced mouse primary cortical neurons with guides for genes of interest (i.e. Cul2 and Vhl, Elob and Vhl, or Cul2 and Lrriq1) and assessed the presence of HIF-1 in cells with an antibody specific for the alpha subunit Hif1a **(Fig. 1c)**. As a positive control, we expressed a stable, non-degradable HIF-1 alpha subunit variant (Hif1a-S) carrying the three mutations P402A/P577A/N813A ^23^. In unmodified mouse cortical neurons, Hif1a was undetectable (**Fig. 1c, top panel**), as might be expected for a constitutively degraded protein, while a robust nuclear signal was observed in neurons expressing stabilized Hif1a-S **(Fig. 1c, bottom panel)**. Interestingly, we found that sgRNA-mediated ablation of CRL2 components (Cullin-2 and Vhl, Elob and Vhl, or Cullin-2 and Lrriq1) resulted in the detection of nuclear Hif1a in neuronal nuclei, although at lower levels compared to transduced Hif1a-S-expressing neurons **(Fig. 1c, mid panels)**. To determine whether prolyl hydroxylases contribute to this mechanism in neurons, we genetically targeted Egln1 and Egln2 with sgRNAs, and used the prolyl hydroxylase inhibitor Roxadustat ^22,26^ (**Supplementary Fig. S3**). In both cases, we found that inhibition of prolyl hydroxylase function led to stabilization of Hif1a and moderate but significant protection from neurodegeneration after axotomy.

We next investigated whether forced expression of stabilized Hif1a-S in mammalian neurons could phenocopy the neuroprotection associated with reduced activity of the CRL2 complex. To this end, primary DRG neurons were transduced on DIV 1 (day in vitro 1) with lentiviral backbone as a control, wild-type HIF-1 alpha subunit (Hif1a-IRES-EGFP), or stabilized, triple mutant HIF-1 alpha subunit (Hif1a-S-IRES-EGFP), and axotomy was performed 10 days post-transduction (**Fig. 2a** and **2b**). Expression of wild-type Hif1a in DRG neurons provided mild protection (**Fig. 2b, center panel**) or no protection at all **(Supplementary Fig. S4b)**, consistent with constitutive degradation. However, stabilized Hif1a-S resulted in severed neurites remaining structurally intact, with a neurite integrity index (NII) as high as ∼0.7 thirty hours after axotomy (**Fig. 2b, right panel**; and **Supplementary Fig. S4b**). Thus, high levels of active HIF-1 can protect neurons from Sarm1-mediated axon degeneration. It is possible that HIF-1 could exert its neuroprotective effect by downregulating Sarm1 expression. We therefore performed Western blots with mouse cortical neurons and found that Sarm1 was not significantly reduced when knocking out *Cul2* and *Vhl* with sgRNA **(Supplementary Fig. S5a)**, a condition that is mildly protective **(Fig. 1b)**. When overexpressing strongly neuroprotective Hif1a-S (using Hif1a-S-IRES-EGFP; the same construct used in **Fig. 2a** and **2b**), Sarm1 protein was only slightly reduced by Western blot **(Supplementary Fig. S5b**; Sarm1 levels with Hif1a-S were ∼90% of control based on densitometric quantification).

**Fig. 2.**
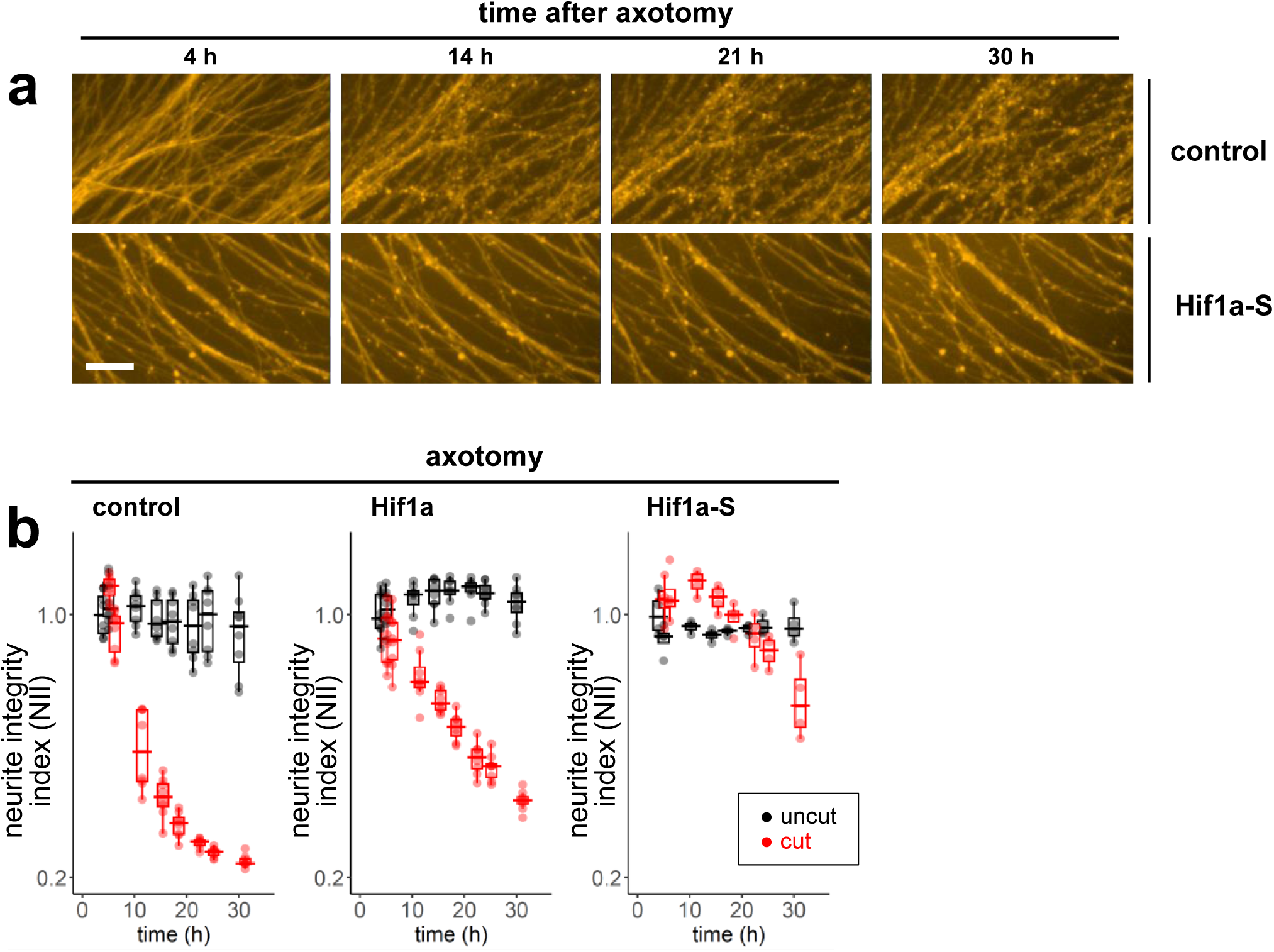
HIF-1 antagonizes Sarm1-mediated neurodegeneration. **a** DRG neurons were spot plated and transduced on DIV 1 with control vector (top row) or degradation-resistant HIF-1 alpha chain (Hif1a-S, bottom row). Axotomy was performed on DIV 11, and neurites distal to the axotomy site were imaged over time. Scale bar, 40 μm. Images are representative of a larger experiment for which the neurite integrity index (NII) was computed and is shown in **(b)**. **b** The neurite integrity index (NII) visualizes axotomy-induced neurodegeneration in DRG neurons transduced with control vector (control, vector backbone including IRES-EGFP, **left panel**), wild-type HIF-1 alpha chain (Hif1a, construct Hif1a-IRES-EGFP, **middle panel**), or degradation-resistant HIF-1 alpha chain (Hif1a-S, construct Hif1a-S-IRES-EGFP, **right panel**). Wild-type Hif1a (**middle panel**) affords some protection, potentially due to saturation of the ubiquitin-dependent Hif1a degradation pathway.

We next used *Drosophila* to determine whether HIF-1-mediated neuroprotection was conserved across humans, rodents, and flies, and whether HIF-1 can protect neurons *in vivo*. The fly genes *sima* and *tango* are the orthologs of mouse HIF-1 alpha and beta subunit genes *Hif1a* and *Arnt*, respectively ^36,37^. Comparison of HIF-1 genes from mouse and fly reveals considerable homologies at the protein level **(Supplementary Fig. 6)** that also encompass DNA-binding and heterodimerization domains ^38^. We expressed mouse Hif1a-S along with a tdTomato reporter in sparse populations of (∼5-10) sensory glutamatergic neurons in the fly L1 wing vein using the MARCM clonal system ^39,40^ (**Fig. 3a**; see methods). We severed axons in the wing vein using microdissection scissors and assayed axon degeneration in axon fibers proximal to the injury site that had been severed from their cell bodies (**Fig. 3a**)(control, intact “bystander” neurons that were not severed are indicated by the “CB” count, **Fig. 3a**). We performed axotomy with flies aged 4 days (5 dpe sample) or 9 days (10 dpe sample), using a control group (expressing UAS-lacZ as a control) and two independent insertions of a UAS-Hif1a-S construct **(Fig. 3b** and **3c)**. Axons were imaged one day after axotomy (1 dpa). Impressively, expression of mouse Hif1a-S conferred significant protection against Wallerian degeneration, with Hif1a-S-expressing severed axons remaining intact for at least a day longer than controls **(Fig. 3b** and **3c)**. Minor differences were observed between Hif1a-S insertion lines, and we note that prolonged expression of 3^rd^ chromosome-inserted Hif1a-S for 9 days (10 dpe) resulted in slight neurotoxicity, suggesting chronically stabilized Hif1a-S can affect cell health. In summary, our data indicate that HIF-1 is normally degraded by prolyl hydroxylases in neurons, but stabilization of HIF-1 can inhibit neurodegeneration driven by dSarm/Sarm1/SARM1 *in vivo*, and this interaction between Sarm1 and HIF-1 signaling is an ancient feature of neurons.

**Fig. 3.**
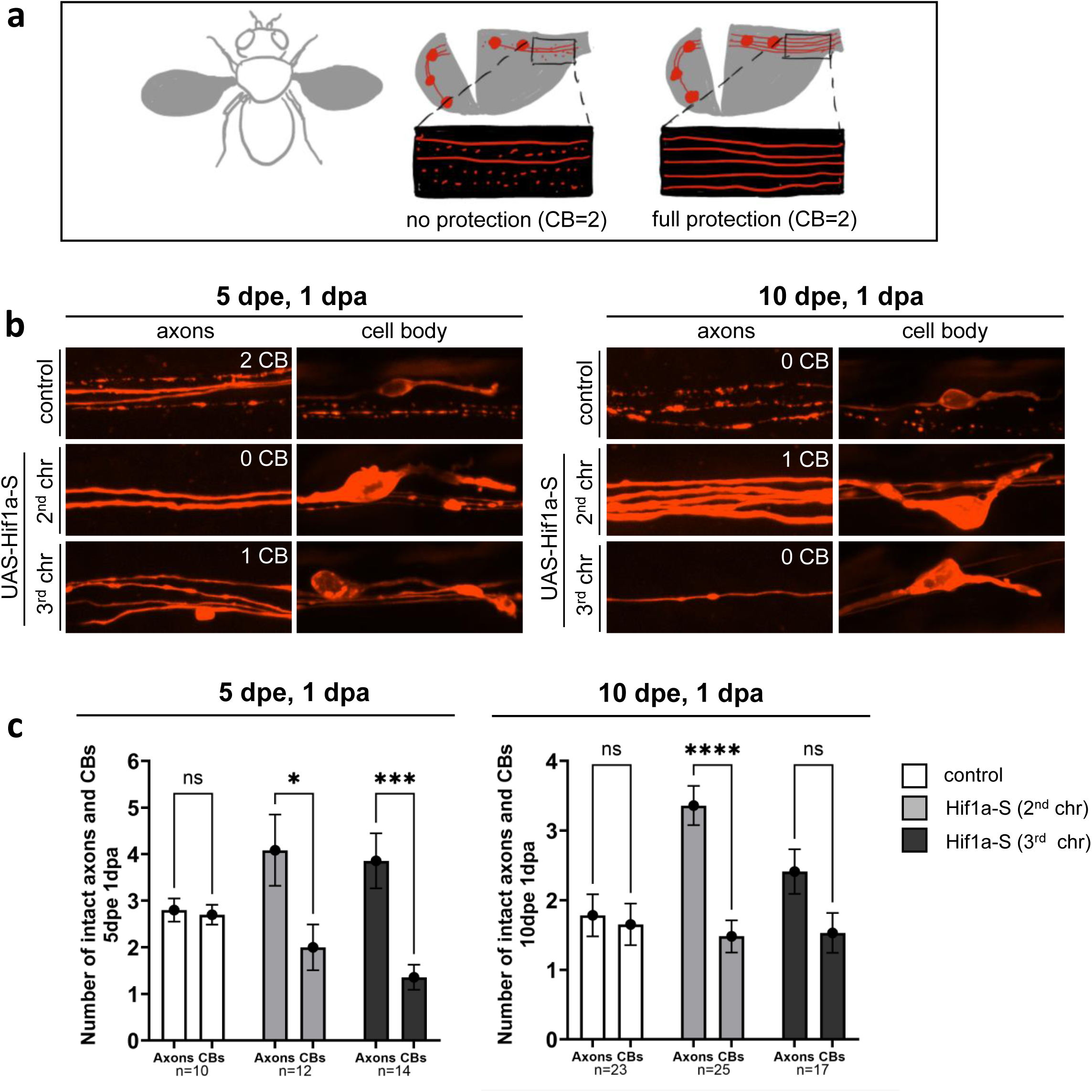
Heterologous expression of stabilized mouse Hif1a-S in fly sensory neurons protects from axotomy-induced neurodegeneration. **a** Schematic representation of wing axotomy and axon imaging. CB (= cell bodies) indicates the number of cell bodies with uncut axons. These axons will remain intact and are not used to evaluate neurodegeneration. **b** Representative images showing protection provided by mouse Hif1a-S in fly sensory neurons. **dpe**, days post eclosion. **dpa**, days post axotomy. **c** Quantifications of intact axons and cell bodies for the experiment show in **(b)**. The results are plotted as mean ± SEM and analyzed using 2way ANOVA with Šídák’s multiple comparison test. **dpe**, days post eclosion. **dpa**, days post axotomy. **n,** number of wings. **\*\*\*\***, p-value < 0.0001. **\*\*\***, p-value 0.0001 to 0.001. **\***, p-value 0.01 to 0.05. **ns**, p-value ≥ 0.05.

### HIF-1 profoundly alters the neuronal transcriptome

To study the transcriptional changes induced by HIF-1-S in primary neurons, we turned to RNA-Seq analysis (**Fig. 4** and **5**). Mouse cortical and DRG neurons were transduced with Hif1a-S or control vector, nuclei were sorted by FACS, and poly(A)-tailed RNA sequences were analyzed with RNA-Seq (see methods for details). Extracted RNA-Seq results are provided in spreadsheet format in Supplementary File ‘RNA-Seq’, and a web-based tool for exploratory analysis of extracted RNA-Seq data can be accessed under https://d3259ptldlzjvg.cloudfront.net/ (currently password-protected; access credentials can be provided upon request). For quantitative evaluation of RNA-Seq results, changes in expression profiles were visualized using volcano plots (**Fig. 4a, left**) and differential expression plots (**Fig. 4a, right**). The number of up- and down-regulated genes (>2-fold up or >2-fold down) ranged in the hundreds, consistent with expectations for HIF-1 transcription factor activity. A significant number of genes were co-regulated across both neuronal cell types assayed (**Fig. 4b**), suggesting that functionally relevant genes for HIF-1-mediated neuroprotection are likely to be found within these co-regulated subsets. A breakdown of RNA-Seq results into different gene types **(Fig. 4c)** revealed the majority of differentially regulated genes were protein-coding. A small subset of lncRNAs (long non-coding RNAs) was also co-regulated in mouse cortical and DRG neurons (**Fig. 4c**). However, the detection of lncRNAs was likely incomplete because RNA-Seq analysis was limited to polyadenylated RNA, and not all lncRNAs have poly(A) tails ^27^.

**Fig. 4.**
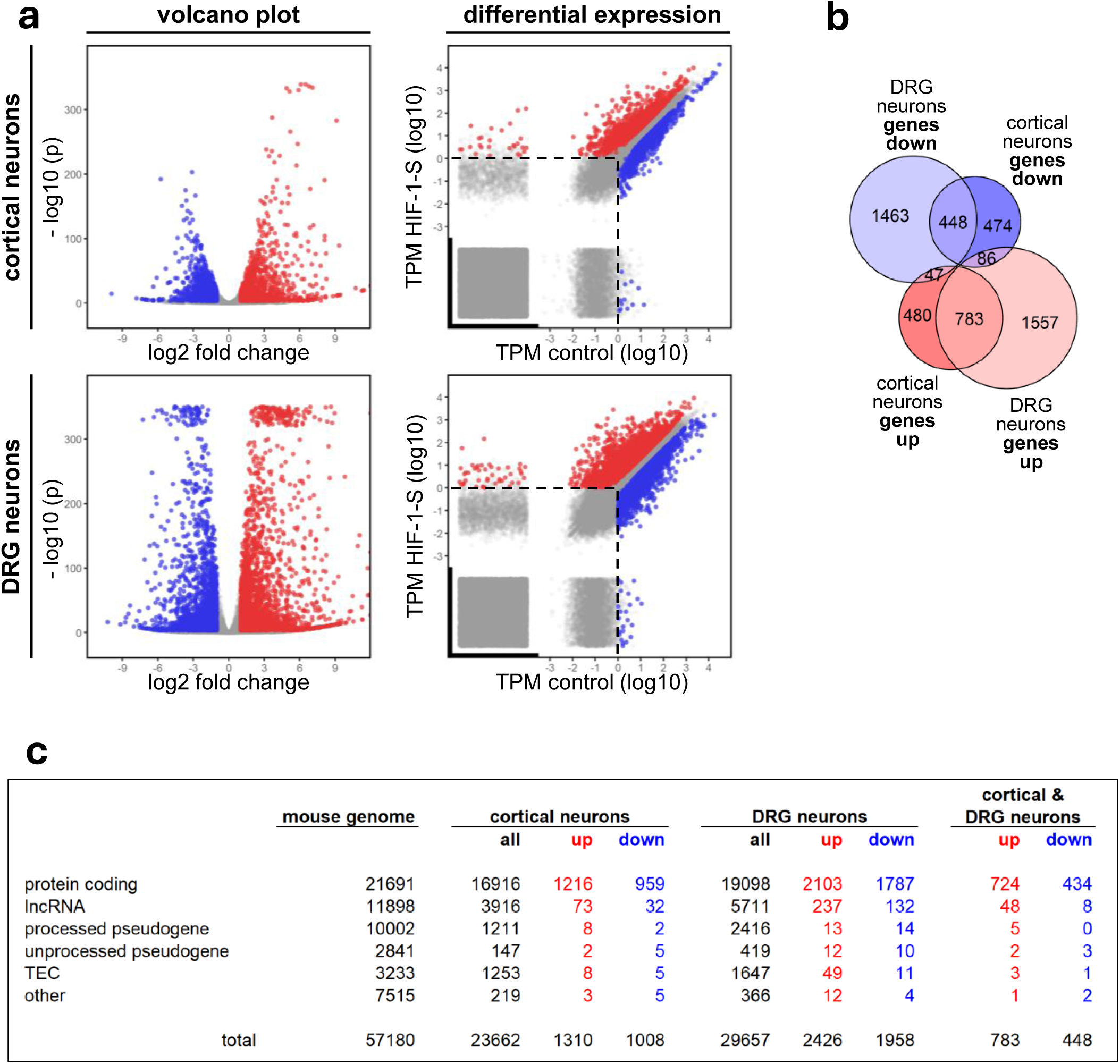
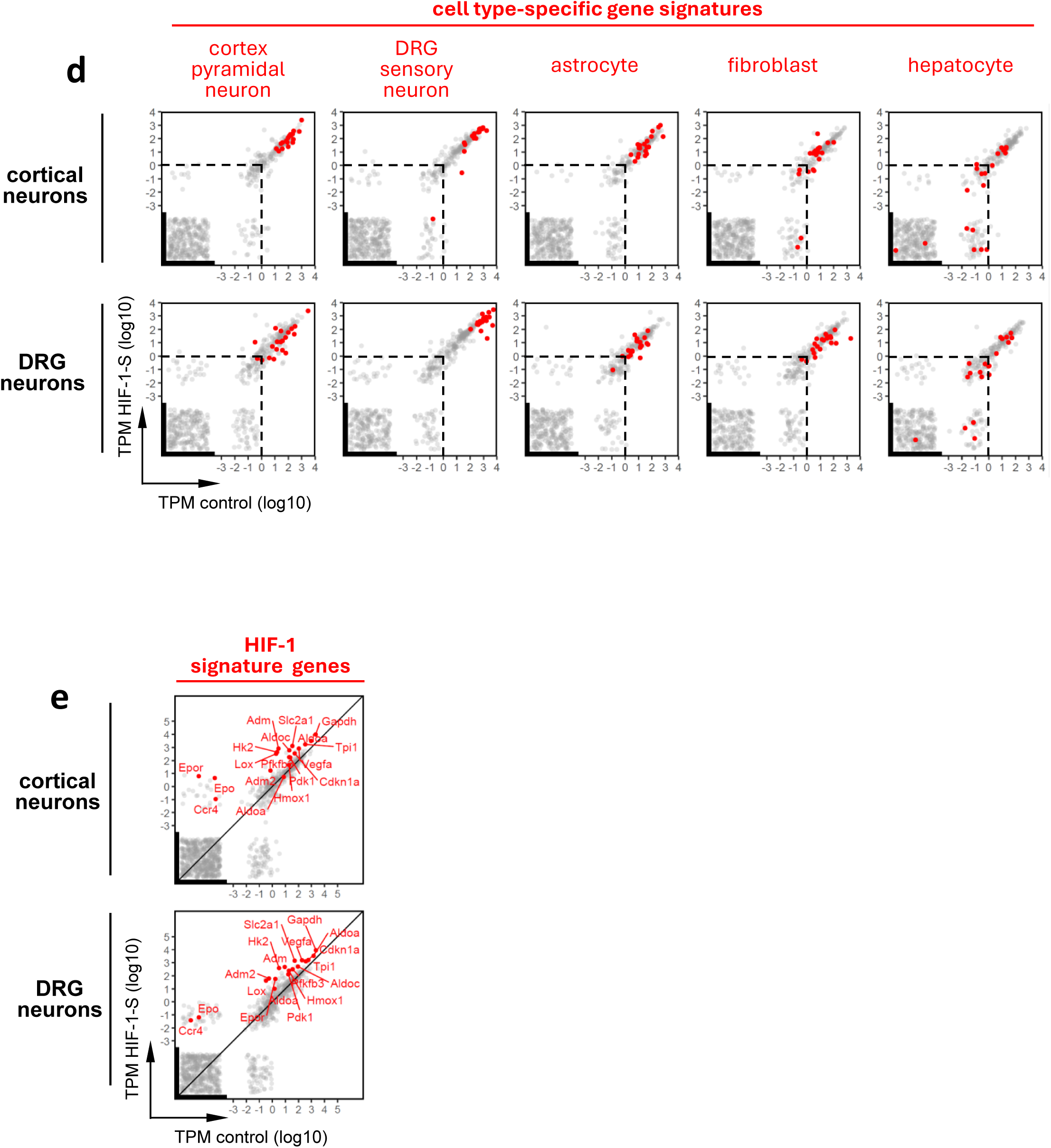
HIF-1-mediated alteration of the neuronal transcriptome a Left panels: Volcano plots showing the dynamic change in gene expression induced by HIF-1-S (x-axis, log2 fold change) and the statistical significance of the observed change (y-axis, - log10 (p)). **Right panels:** differential expression plots with normalized expression values (TPM = transcripts per million) for each gene in control condition (x-axis, TPM control) and experimental condition with HIF-1-S (y-axis, TPM HIF-1-S). Also included in the differential expression plots are genes that are not expressed in the analyzed cells (TPM = 0); these genes are shown over the thickened axis portions, at randomized positions. Dashed line, estimated threshold for biologically meaningful expression at TPM = 1. Red, genes upregulated 2-fold or higher; blue, genes downregulated 2-fold or lower; gray, remaining genes **b** Euler diagram visualizing overlapping groups of HIF-1-regulated genes (including all types of genes), comparing cortical neurons and DRG neurons. Criteria: TPM > 1, log2 fold change > 1.0 (up) or <-1.0 (down), FDR-adjusted p value < 0.001. **c** Breakdown of gene numbers by cell type and gene type, based on the transcriptome used for alignment (from ensembl.org), which lists a total of 57,180 genes with multiple categories. lncRNA = long non-coding RNA, TEC = genes “to be experimentally confirmed”. **d Gene signatures show collected RNA-Seq data are highly specific for cortical and DRG neurons.** Gene signatures obtained from single-cell RNA-Seq datasets (from cellxgene.cziscience.com), using the 20 highest ranking cell type-specific marker genes, were projected onto RNA-Seq differential expression plots. Dotted lines mark the common threshold estimate for biologically relevant TPM values (at TPM = 1). See supplement “gene_sets”, tab “cell type signatures” for details. **e HIF-1-S induces expression of known HIF-1 target genes in neurons.** See supplement “gene_sets”, tab “HIF-1-induced genes” for details. **d,e** Unexpressed genes (TPM = 0) are shown over thickened axis portions with randomized position.

**Fig. 5.**
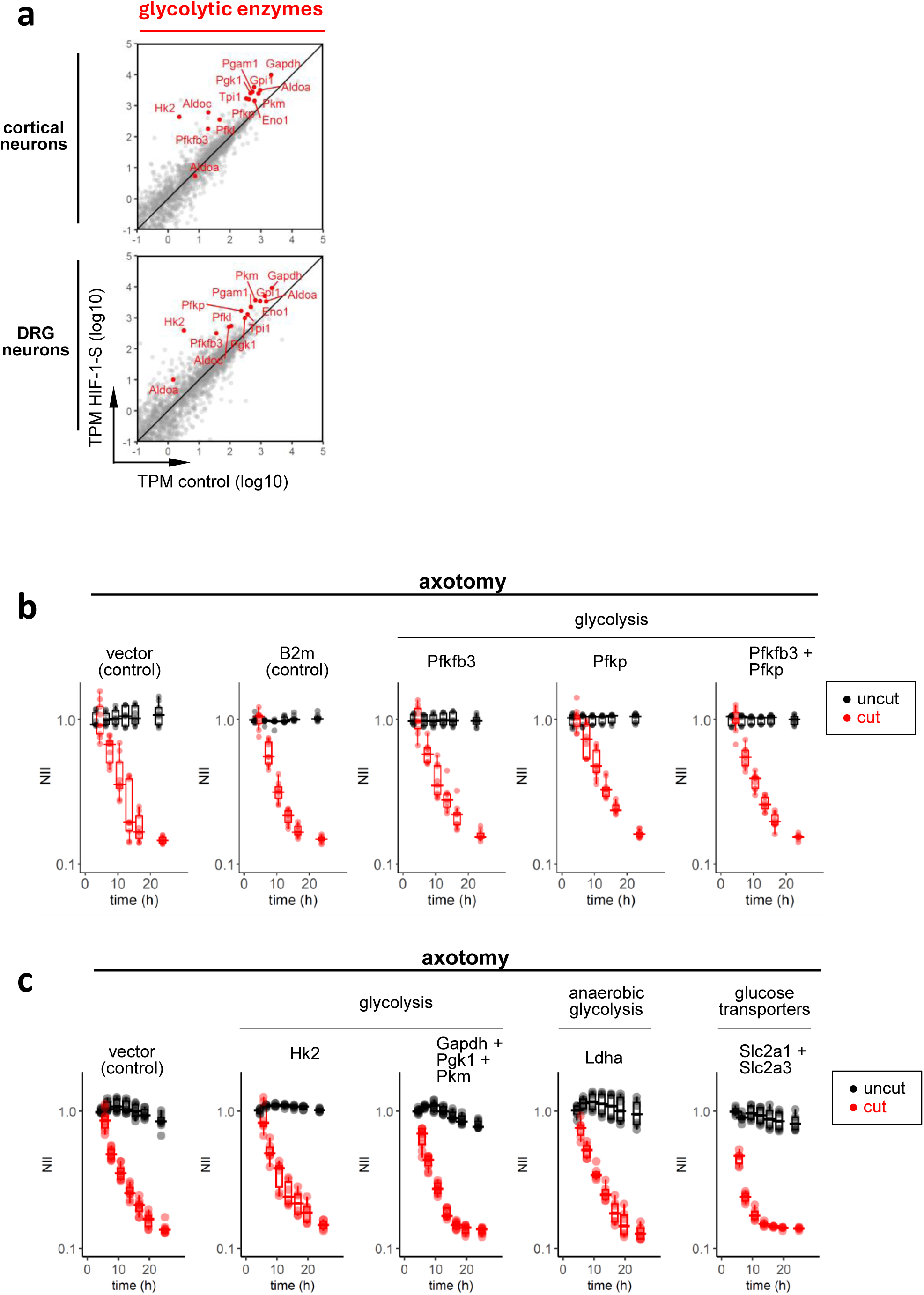
HIF-1-S potently induces transcription of glycolytic genes, but overexpression of glycolytic enzymes does not result in neuroprotection. **a** HIF-1-S-induces all 10 classes of glycolytic enzymes in mouse cortical and DRG neurons (see Supplementary File “gene_sets”, tab “glycolysis”; and Supplementary Table T5) **b,c** Axotomy of DRG neurons transduced with control vector or expression constructs. **b** B2m, beta-2 macroglobulin; Pfkfb3, 6-phosphofructo-2-kinase; Pfkp, phosphofructokinase, platelet type. Pfkfb3 is an allosteric activator of the metabolically active posphofructokinase complex, a tetrameric combination of Pfkp, Pfkl, or Pfkm. **c** Hk2, hexokinase; Gapdh, glyceraldhyde-3-phosphate dehydrogenase; Pgk1, phosphoglycerate kinase; Pkm, pyruvate kinase; Ldha, lactate dehydrogenase; Slc2a1, glucose transporter 1 (GLUT1); Slc2a3, glucose transporter 3 (GLUT3).

To verify that the obtained RNA-Seq data were specific for neurons, we used gene signatures compiled from publicly available single-cell RNA-Seq datasets (https://cellxgene.cziscience.com/) **(Supplementary File “gene_sets”, tab “cell type signatures”)**. Projecting signatures for pyramidal neurons, sensory neurons, and non-neuronal cell types onto differential expression plots provided clear evidence that our data represented mouse cortical and DRG neurons (**Fig. 4d**). Examination of a panel of known HIF-1-induced target genes **(Supplementary File “gene_sets”, tab “HIF-1-induced genes”)** ^28,29^ suggested that Hif1a-S activity in neurons recapitulated HIF-1 functionalities discovered in other cell types, as almost all HIF-1 signature genes were found to be upregulated in neurons **(Fig. 4e)**. Hif1a-S even boosted genes that are normally not expressed in neurons, and are unrelated to neuron function, such as *Epo* (erythropoietin), *Ccr4* (chemokine receptor 4), *Vegfa* (VEGF), and *Adm* (adrenomedullin).

Examining RNA-Seq results for functional clues that could explain HIF-1-mediated neuroprotection, we first looked at enzymes involved in NAD^+^ biosynthesis and NAD^+^ consumption. As for NAD^+^ biosynthesis, a comprehensive list of 15 enzymes **(Supplementary File “gene_sets”, tab “NAD^+^ biosynthesis”)** in this pathway ^30^ was not changed in a meaningful way by HIF-1-S (**Supplementary Table T1**). *Nmnat2*, the gene encoding the central enzyme for NAD^+^ synthesis, was slightly reduced, and there was weak enhancement of expression of *Ido2* (indoleamine-2,3-dioxygenase, involved in de novo NAD^+^ synthesis). NAD^+^ consumption has been mainly ascribed to two major enzyme families, i.e. PARPs and sirtuins. For both, we did not find evidence for strong HIF-1-mediated dynamic regulation (**Supplementary Table T2**). Genes for NAD^+^ consuming enzymes Cd38 (ADP-ribosyl cyclase/cyclic ADP-ribose hydrolase 1) and Bst1 (CD157, ADP-ribosyl cyclase/cyclic ADP-ribose hydrolase 2) were expressed below meaningful levels in mouse neurons and unchanged by HIF-1-S. In aggregate, transcriptome analysis suggests that NAD^+^ production and consumption are not significantly altered in neurons by HIF-1-S.

As both NAD^+^ and ATP are massively depleted after Sarm1 activation, we examined effects on mitochondrial pathways of energy metabolism **(Supplementary File “gene_sets”, tab “energy”)**, i.e. the electron transport chain (complexes I-IV) and ATP synthase (complex V) (**Supplementary Fig. S7a**), citric acid cycle, and pyruvate metabolism (**Supplementary Fig. S7b**). In these pathways, our RNA-Seq results showed only minor or punctual HIF-1-S-induced changes. Noteworthy is the upregulation of mitochondrial complex IV component Cox4i2 (**Supplementary Table T3**). It has been proposed ^31^ that a HIF-1-mediated switch from complex IV subunit Cox4i1 to Cox4i2, along with upregulation of protease Lonp1, makes mitochondrial respiration more efficient. Our RNA-Seq results reported this exact pattern – reduction of Cox4i1, strong upregulation of Cox4i2, and induction of Lonp1 – for both investigated neuronal cell types. More than other cells, neurons rely on the pentose phosphate pathway for supplying antioxidant capacities in the form of reduced glutathione ^32^. Enzymes in this pathway were affected by HIF-1 only to a minor degree (**Supplementary Table T4**).

The glycolysis pathway experienced a strong transcriptional enrichment in Hif1a-S-expressing neurons. In each of the 10 canonical enzyme groups of glycolysis, at least one enzyme was found to be strongly induced (**Fig. 5a; Supplementary File “gene_sets”, tab “glycolysis”**; and **Supplementary Table T5**). Other glucose-related genes with strong induction were glucose transporters *Slc2a1* (GLUT1, glucose transporter 1) and *Slc2a3* (GLUT3, glucose transporter 3), as well as *Ldha* (lactate dehydrogenase), a central enzyme in anaerobic glycolysis (**Supplementary Table T5**). To test the simple hypothesis that enhanced glycolysis may result in significant neuroprotection, we expressed various glycolytic enzymes in DRG neurons, individually or in combination, and quantified axotomy-induced neurodegeneration over time (**Fig. 5b** and **5c**). Neurodegeneration was found to be unaffected in these experiments, which included overexpression of *Pfkfb3* (6-phosphofructo-2-kinase), a central regulator of glycolytic flux. Overexpression of lactate dehydrogenase (Ldha) and of glucose transporters (Slc2a1 and Slc2a3) also failed to protect neurons **(Fig. 5c)**. It is possible that many more, or all glycolytic enzymes in concert, must be upregulated to achieve significant neuroprotection.

### Sustained HIF-1 expression and interaction with DNA are required for neuroprotection

Lacking a clear transcriptional change that might explain the neuroprotection by Hif1a-S, we sought evidence that its neuroprotective effects were through gene regulation. Hif1a is the alpha chain of the heterodimeric transcription factor HIF-1 and pairs with the beta chain Arnt. The N-termini of both alpha and beta subunits of HIF-1 interact with DNA, with bHLH domains at the N-termini recognizing short motifs in the genome known as hypoxia response elements (HREs) ^33,34^. To test whether HIF-1 binding to HREs and transcriptional activity is essential for its neuroprotective effect, we generated two HIF-1 alpha chain constructs lacking a major portion of the 70aa N-terminal bHLH DNA binding domain: Δ2-28 Hif1a-S (i.e. preserving the stabilizing mutations P402A/P577A/N813A, but introducing a defect in the bHLH domain), and a dominant-negative Hif1a fragment (aa 30-389, Hif1a^DN^). Primary DRG neurons were transduced with control vector, Δ2-28 Hif1a-S, or Hif1a^DN^, and axotomy was performed 10 days after transduction. We found the expression of Δ2-28 Hif1a-S did not provide any protection against Wallerian degeneration **(Fig. 6a)**, demonstrating that Hif1a-S requires DNA binding for its neuroprotective effect. A similar result was obtained with Hif1a^DN^ (**Fig. 6b**). These experiments support the notion that DNA binding and transcriptional activity of HIF-1 is crucial to protect severed neurites.

**Fig. 6.**
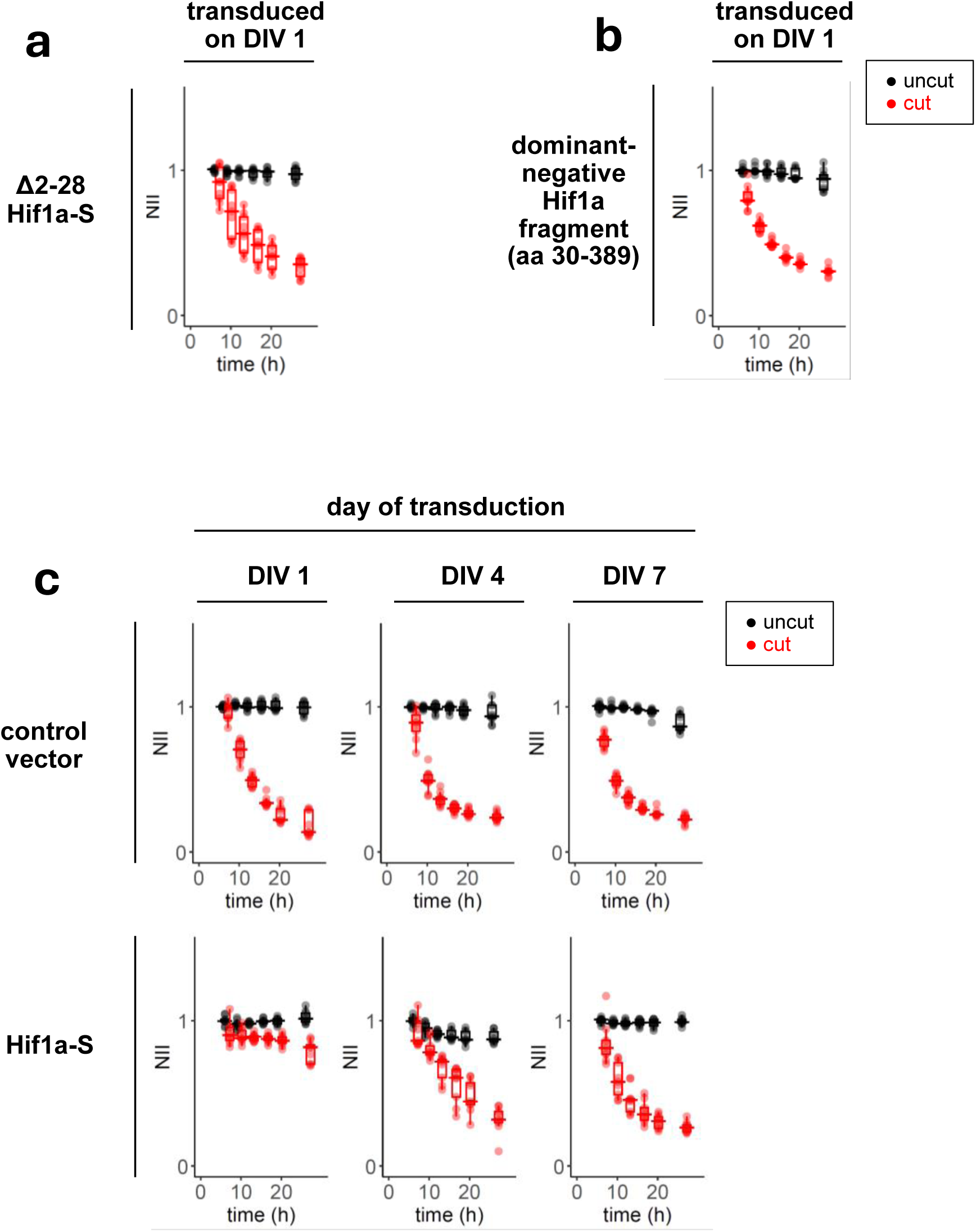
Sustained HIF-1 expression and interaction of HIF-1 with DNA is required for neuroprotection. **a** Axotomy of DRG neurons transduced with Δ2-28 Hif1a-S, which lacks part of the N-terminal DNA interaction domain but retains stabilizing mutations P402A/P577A/N813A. **b** Axotomy of DRG neurons transduced with a dominant-negative Hif1a fragment (Hif1a aa 30-389). **c** Mouse DRG neurons were transduced on DIV 1, DIV 4, or DIV 7 with degradation-resistant Hif1a-S, or control vector. Axotomy for all conditions was performed on DIV 11, and neurodegeneration over time was quantified using the neurite integrity index (NII). **a,b,c** All panels report data from the same experiment. The control transduction on DIV 1 in **(c)** serves as control reference for **(a)** and **(b)**.

If HIF-1-mediated neuroprotection required transcription, this might occur over a slow time scale. We asked how long HIF-1 needed to be present in neurons to protect against axotomy. DRG neuron cultures were transduced with control vector or Hif1a-S on DIV 1 (as in our standard protocol), but also on DIV 4 and on DIV 7. Performing axotomy on DIV 11 on all samples, we observed that delivery of Hif1a-S on DIV 1 produced maximum protection (**Fig. 6c, left panels**), while its delivery on DIV 4 was far less effective (**Fig. 6c, center panels**), and delivery on DIV 7 had no effect. We used mCherry as a control reporter for general translational activity, and EGFP as a direct reporter for Hif1a-S (i.e. mCherry was delivered with a separate construct, while Hif1a-S and EGFP were jointly expressed in a bicistronic cassette, both driven by the human Synapsin 1 promoter), since either could be affected by Hif1a-S. On DIV 11 (i.e. on axotomy day), the resulting mCherry signal was similar for all transduction groups, and Hif1a-S reporter EGFP was equally expressed for transductions on DIV 1 and DIV 4 but dropped afterwards. In summary, these data are consistent with the notion that Hif1a-S must be present in neurons for days for its neuroprotective effect and are consistent with the notion that Hif1a-S suppresses Sarm1-mediated neurodegeneration by altering gene expression in neurons.

### HIF-1 blocks neurodegeneration independent of neuron type and Sarm1 activation mode

The mechanism by which HIF-1 interacts with Sarm1 can be explored using tools that activate Sarm1 differentially. We first used Vacor, a neurotoxin that directly activates endogenous Sarm1 by interacting with its autoinhibitory ARM domain ^10^. Mouse cortical neurons were transduced on DIV 1 with control lentivirus or Hif1a-S. On DIV 11, cells were exposed to various concentrations of Vacor (**Fig. 7a**). Hif1a-S conferred robust protection against Vacor-induced Sarm1 activation and neurodegeneration, keeping cell bodies and neurites morphologically healthy for at least 22 hours with 10 μM Vacor. In contrast, control neurons exhibited rapid degradation of cell bodies and neurites within 2-3 hrs after Vacor exposure (**Fig. 7a**).

**Fig. 7.**
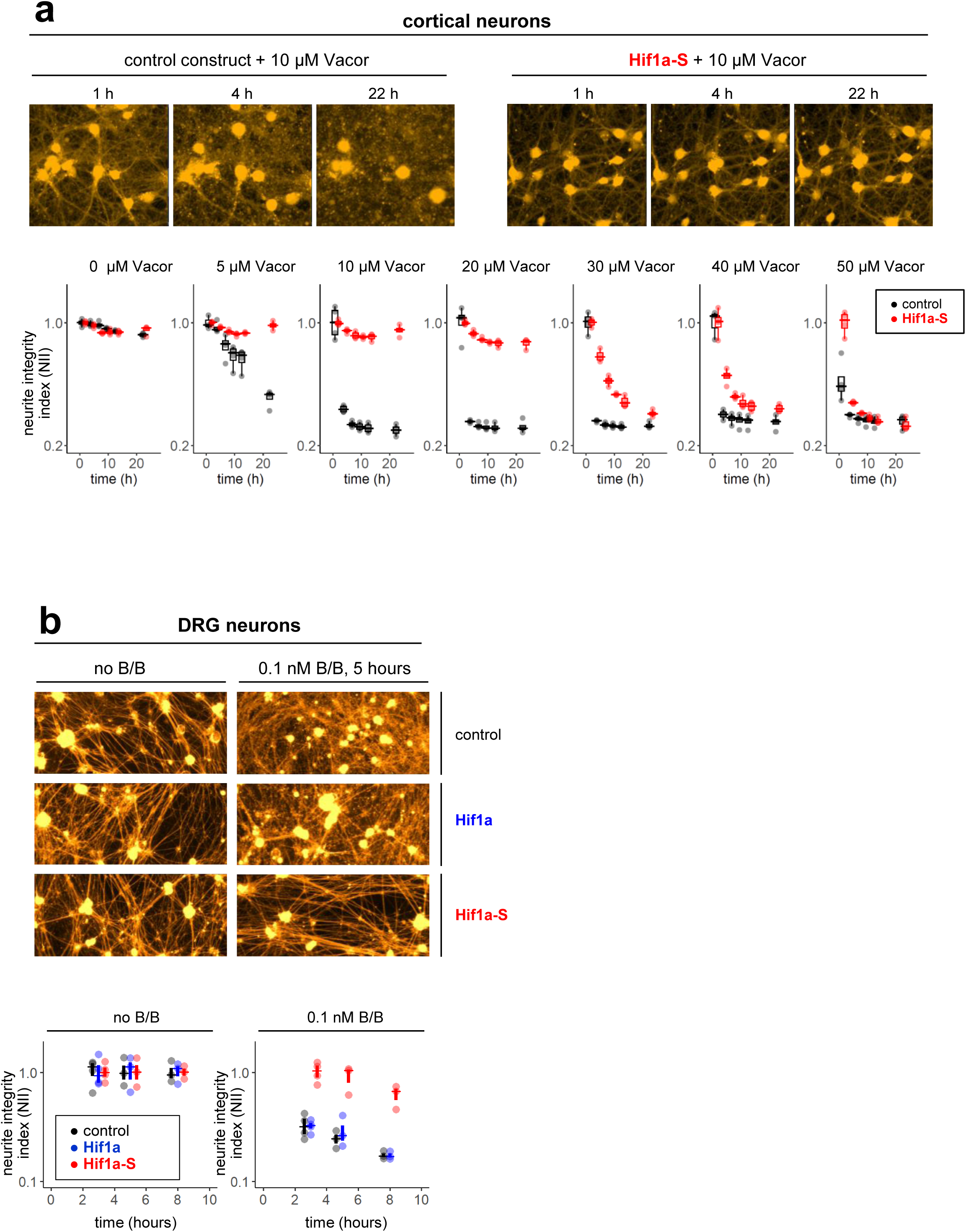
HIF-1 protects against Sarm1 in different neuronal cell types and with different Sarm1 activation modes. **a** Mouse cortical neurons were transduced on DIV 1 with control vector or degradation-resistant Hif1a-S. On DIV 11, Vacor, a chemical activator of Sarm1, was added, and cells were imaged for several hours. **Top panels**, representative excerpts from images for select conditions. **Bottom panel**, neurite integrity index (NII) time course for all experimental conditions. **b** Sparsely plated mouse DRG neurons were transduced on DIV 1 with control construct, wild-type Hif1a, or degradation-resistant Hif1a-S. All conditions also received Fh-TIR (fused to mCherry), which contains the Sarm1 NADase domain (TIR domain) and can be activated by the homodimerizer B/B, which was added on DIV 11. Cells with and without B/B homodimerizer were imaged over time. **Top panels,** representative images. **Bottom panels,** neurite integrity index (NII) computed for all experimental conditions.

The TIR domain of SARM1 possesses intrinsic NAD^+^ hydrolase activity, which is activated upon homodimerization, and this leads to depletion of NAD^+^ and ATP to catastrophically low levels, and cell degeneration ^17^. We generated a fusion protein, Fh-TIR-mCherry, which combined mCherry with the SARM1 TIR domain and the homodimerization domain FKBP(F36V). Fh-TIR is inactive as a monomer but exhibits NAD^+^ hydrolase activity upon dimerization by the small molecule B/B (AP20187) ^17,35^. Sparsely plated mouse DRG neurons were transduced with control vector, wild-type Hif1a, or stabilized Hif1a-S. Neurons were also transduced with Fh-TIR-mCherry, and its expression levels were indistinguishable in all conditions. Fh-TIR dimerization was induced by addition of the B/B compound on DIV 12 (**Fig. 7b**; and **Supplementary Fig. 8)**. Control as well as wild-type Hif1a-expressing neurons underwent degeneration 5 hours after Fh-TIR dimerization. In contrast, stable Hif1a-S resulted in significant neuroprotection, with neurites showing intact gross morphology even 8 hours after the B/B application. This observation further supports the notion that stabilized Hif1a-S protects neurons from SARM1-mediated neurodegeneration, even when Sarm1 activity is narrowed to its TIR domain-mediated NAD^+^ hydrolase function.

### HIF-1-S blocks loss of NAD^+^ after Sarm1 activation

A key pro-degenerative activity of Sarm1 is depletion of NAD^+^. To determine whether Hif1a-S mitigated the ability of Sarm1 to deplete NAD^+^ in neurons, we assayed Sarm1 activation after Vacor addition. We expressed Hif1a-S in cultured mouse cortical neurons, activated Sarm1 with Vacor, and measured NAD^+^ and its reduced counterpart NADH over time using a bioluminescence-based assay (**Fig. 8a**). Impressively, Hif1a-S effectively inhibited the loss of NAD^+^ after Sarm1 activation. At 5 μM Vacor, cells with Hif1a-S maintained stable NAD^+^ and NADH levels, in contrast to unprotected cells, which exhibited a sharp decline in both metabolites **(Fig. 8a, left panels)**. Even with 10 μM Vacor, which induces stronger Sarm1 activation, Hif1a-S stabilized NAD^+^ levels, albeit to a lesser extent **(Fig. 8a, right panels)**. In parallel, we quantified neurodegeneration over time for the same batch of cells **(Fig. 8b)**. At 5 μM Vacor, all cells remained morphologically intact for 6 hours, regardless of Hif1a-S protection. At 10 μM Vacor, unprotected cells exhibited visible degeneration after 6 hours. Notably, Hif1a-S-transduced cells maintained morphological integrity even 48 hours after Vacor addition **(Fig. 8b, right panels)**, despite experiencing mild NAD^+^ reduction at the 6-hour mark **(Fig. 8a, right panels)**. Given that Vacor acts directly on Sarm1, these data argue that gene expression changes that occur in Hif1a-S-expressing neurons can suppress the ability of the Sarm1 hydrolase to drive NAD^+^ depletion in neurons.

**Fig. 8.**
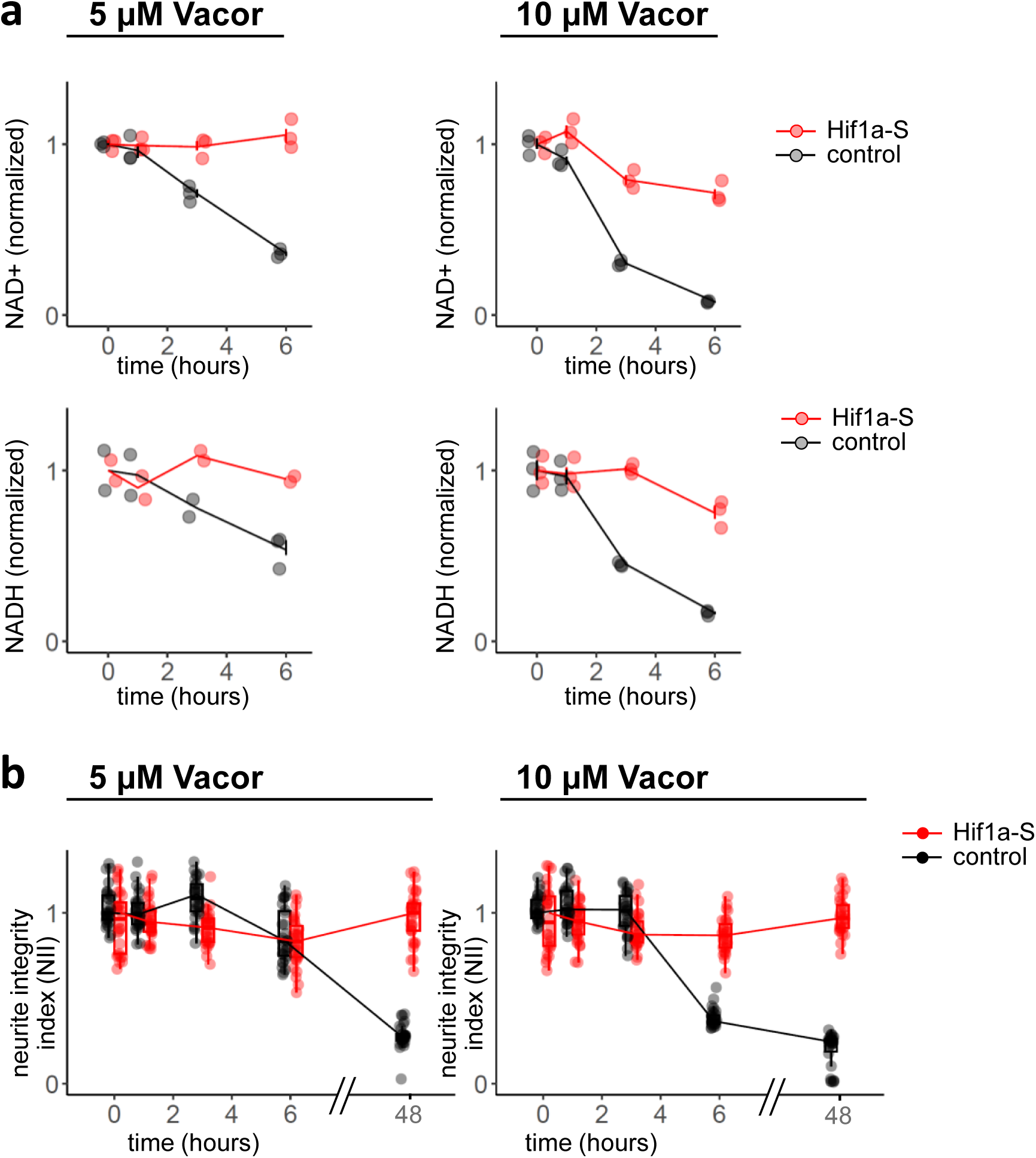
HIF-1 prevents precipitous loss of NAD^+^ and NADH after Sarm1 activation. **a** Mouse cortical neurons were transduced on DIV 1 with stabilized HIF-1 alpha chain (Hif1a-S) or empty vector (control). On DIV 11, Vacor was added at a concentration of 5 μM (left panels) or 10 μM (right panels). NAD+ and NADH were measured with colorimetric assay after different periods of Vacor exposure. Counts for NAD+ and NADH were normalized and plotted against time. **b** In parallel to NAD+ analysis, for the same batch of mouse cortical neurons shown in (a), the neurite integrity index (NII) was determined. Cells were imaged at the same time points chosen for NAD+ and NADH measurement (0, 1, 3, and 6 hours), plus an additional late time point at 48 hours after adding Vacor. Intact cell morphology at 48 hours indicates that Hif1a-S can help cells to survive periods of NAD+ depression.

## Discussion

After nervous system injury or in neurodegenerative disease, neurons are forced to confront a complex array of challenges to survive and remain functionally integrated into circuits. Physical injury can directly damage the vasculature, and vascular defects are common in neurodevelopmental disorders and neurodegenerative diseases, including multiple sclerosis ^41^, Parkinson’s, Alzheimer’s and Huntington’s disease ^42^. Neurons that are damaged beyond repair will likely degenerate, often in a Sarm1-dependent fashion ^43^. However, neurons that are not directly damaged (bystanders) can temporarily modify their function but later recover and survive ^44,45^. This is impressive since tissue damage, inflammation, or degeneration can also result in dramatic changes in vascular function that deprives neurons of nutrient delivery and efficient gas exchange. In this study, we made the unexpected discovery that Sarm1 signaling can be directly mitigated by the O_2_-sensitive transcription factor HIF-1. Under normoxic conditions, HIF-1 is degraded after prolyl hydroxylases use O_2_ to hydroxylate the HIF-1 alpha subunit (Hif1a) for recognition by Vhl and delivery to the proteasome ^16^. Manipulations that lead to HIF-1 stabilization, including loss of Cullin-2 RING E3 ligase (CRL2) components, or mutations of key hydroxylation sites in Hif1a, suppressed the ability of activated Sarm1 to drive degeneration of axons (and cell bodies) after axotomy. These findings support the view that hypoxia can meaningfully regulate the pro-degenerative function of Sarm1. We propose that HIF-1 mitigates the pro-degenerative function of Sarm1 through broad changes in transcription, enhances the ability of neurons to deal with stress during responses to brain injury or in neurodegenerative disease, and increases their chances to survive and remain functionally intact.

We used genome-wide sgRNA screens in human cell lines to find molecules required for constitutively active SARM1 (SARM1ΔARM) to drive neurodegeneration. We undertook extensive efforts to optimize our screening system, which included measures to guarantee correct integration of SARM1ΔARM and its stable expression along with sgRNAs, as well as screening several cell lines to maximize screening efficiency. Ultimately, only a handful of hits resulting from cell line-based, genome-wide sgRNA screens could be validated in mouse primary neurons, where their guide-mediated knockdown conferred moderate protection against axotomy (e.g. Cullin-2, Vhl, Lrriq1, and Fem1 family proteins). We propose that few genes exist that, when ablated genetically as single loss-of-function mutants, can suppress Sarm1-mediated neurodegeneration. Individual loss of Cullin-2 or of CRL2 receptor Vhl only weakly suppressed activated Sarm1 in primary cells, but simultaneous knockdown of Cullin-2 with associated receptors (Vhl, Lrriq1, or Fem1 family proteins) or canonical adaptors (Elob, Eloc) resulted in stronger suppression. Consistent with their acting through HIF-1 and its regulation by O_2_-sensing prolyl hydroxylases, overexpression of wild type HIF-1 alpha subunit (Hif1a) resulted in moderate to no suppression of neurodegeneration after axotomy, presumably due to its constitutive degradation in normoxia. In contrast, expression of a stabilized HIF-1 alpha subunit variant (Hif1a-S) strongly suppressed the pro-degenerative effects of activated Sarm1. Additional support for a role for O_2_ in HIF-1-mediated regulation of Sarm1 comes from our manipulations of prolyl hydroxylases, using genetic and pharmacological methods. In these experiments, we observed weak but significant suppression of neurodegeneration with guide-mediated ablation of prolyl hydroxylase genes *Egln1* and *Egln2*, and using the prolyl hydroxylase inhibitor Roxadustat.

Transcriptome analysis of primary neurons (DRGs and cortical neurons) expressing Hif1a-S revealed a significant change in neuronal transcription that affected hundreds of genes, including a set of known HIF-1 targets. We found no evidence for transcriptional changes in NAD^+^ synthesis pathways, or gene families related to NAD^+^ consumption. Mitochondrial energy pathways were altered in very limited ways, and the significance of a few differentially regulated genes in these pathways must ultimately be investigated experimentally. However, there was a marked induction of transcripts required for glycolysis, which could function to counteract the drop in ATP that follows Sarm1 activation ^35^, although we found that expression of individual glycolytic genes, or combinations thereof, was not sufficient for neuroprotection. Our data are consistent with the notion that HIF-1 counteracts Sarm1 via transcription, because Hif1a-S had to be expressed for several days to impart resistance to neurodegeneration and this effect was dependent on an intact DNA binding domain. Since HIF-1 is a transcription factor, its mechanism is most likely manifest in the transcriptional changes reported by our RNA-Seq datasets, which we publish here in full and accessible formats, including a freely available web tool for exploration. Several hundred HIF-1 target genes are co-regulated in mouse cortical and DRG neurons and are likely to include the subset functionally relevant for neuroprotection. Precisely how HIF-1 exerts these neuroprotective effects is an exciting question for the future, and if fully understood, might be used to further enhance anti-Sarm1 therapeutic approaches in neurodegeneration that target Sarm1 hydrolase function.

We envision at least two models for how HIF-1 suppresses Sarm1-mediated neurodegeneration. First, HIF-1 could upregulate pathways that (through mechanisms we are unable to resolve by gene expression analysis) enhance NAD^+^ production or slow NAD^+^ consumption, which would counteract the drop in NAD^+^ following Sarm1 activation. Alternatively, HIF-1 could induce expression of a factor in cells that can inhibit Sarm1 signaling directly (through physical interaction with Sarm1) or indirectly (through blockade of Sarm1 signaling partners). The observation that HIF1a-S also antagonizes dimerized Fh-TIRa, potent NAD^+^ hydrolase with no other Sarm1 domains, argues that HIF1a-S-mediated changes in cells can directly counteract the hydrolase activity of Sarm1. This could be a factor that directly binds the TIR domain of Sarm1, or it might inhibit a downstream signaling pathway necessary for sustained Sarm1-dependent NAD^+^ consumption in intact cells.

How hypoxia may play a role in regulating Sarm1 function in the intact nervous system is an exciting question, and seems highly relevant to neurodegenerative processes in stroke, traumatic brain injury (TBI), and other neurological diseases. A growing body of work supports the notion that, while acute and severe hypoxia is damaging to the CNS, moderate and even repetitive hypoxia can be neuroprotective ^46^. Indeed, hypoxia was found to extend lifespan and neurological function in a mouse model of rapid aging ^47^, and hypoxia rescued neurological defects (but not cardiovascular problems) in preclinical models of Friedreich’s ataxia ^48^. Sarm1 is directly regulated by its ability to sense metabolic status through its binding of NAD^+^ and NMN, metabolites that respectively inhibit and activate Sarm1 NAD^+^ hydrolase ^13^. Our work adds hypoxia to the growing list of physiological mechanisms that can impinge on Sarm1 signaling, and we propose that transient suppression of Sarm1 activity by hypoxia could be a mechanism to mitigate its pro-degenerative effects until normal O_2_ levels are restored.

## Methods

### Mice

Wild-type C57BL/6J mice (Jackson stock no. 000664), Sarm1^-/-^ C57BL/6J mice (Jackson stock no. 018069), and C57BL/6J mice constitutively expressing Cas9 (Jackson stock no. 028239) ^51^ were purchased from The Jackson Laboratory (Bar Harbor, ME). Neurons from Cas9^+/+^ embryos were used for experiments involving Cas9 sgRNA-mediated gene knockdown, while cells from both wild-type and Cas9^+/+^ embryos were used for overexpression experiments. Mouse breeding and handling was in accordance with OHSU IACUC and DCM institutional guidelines and protocols.

Timed pregnant mice were made by mating Cas9^+/+^ males and females in pairs for a period of 36-48 hours. Females were weighed every other day to check for pregnancies. DRGs were obtained from E13 or E14 embryos ^52^, and mouse cortical neuron cultures were set up from E15 or E16 embryonic brains ^53,54^.

### Constructs

Retroviral vector Cas9 in pQCXIH was made by subcloning hSpCas9 (human codon-optimized S. pyogenes Cas9) from lentiCas9-Blast (Addgene plasmid #52962) into plasmid pQCXIH (Clontech).

For guide expression in mouse primary neurons, we modified vector lentiGuide-Puro (Addgene plasmid #52963). For single guide expression, we created lentiGuide-P.Syn-EGFP by swapping the puromycin cassette in lentiGuide-Puro with EGFP driven by the human Synapsin 1 promoter, for robust and neuron specific EGFP reporter expression. For simultaneous delivery of two guides, we engineered lentiGuideDuo-P.Syn-EGFP, with two sgRNA expression cassettes driven respectively by the human U6 and the human H1 promoter, arranged in a converging configuration, as previously described ^55^. In pilot experiments, we verified robust editing of two simultaneously targeted genes with lentiGuideDuo-P.Syn-EGFP. sgRNA constructs were made using guide sequences from the Brunello human sgRNA library (Addgene #73178 ^18^, 4 guides per gene); the Brie mouse sgRNA library (Addgene #73633 ^18^, 4 guides per gene); and the Gouda mouse sgRNA library (Addgene #136987 ^56^, 2 guides per gene).For overexpression experiments, a vector of the pLenti series was engineered to drive transgene expression with the human Synapsin 1 promoter, and named pLenti-P.Syn. Construct maps and sequences are available upon request.

### Lentivirus production

HEK293T cells were plated 24 h before transfection at 7-10 x 10^6^ cells in 9 ml D10 medium per 10 cm dish. The transfection mix for a 10 cm dish included: 6 ug transfer vector, 4 ug packaging vector (psPAX2 for lentivirus, pCG gag/pol for retrovirus), 2 ug envelope vector (pMD2.G), 1.5 ml Opti-MEM I, and 36 µl TransIT-293 transfection reagent (Mirus MIR 2700). The transfection mix was incubated for 15-30 min at room temperature, then added dropwise to the HEK293T cells, which were transferred to a 37°C, 5% CO2 incubator. 24 h after transfection, all media was aspirated and replaced with 9 ml D10 medium (or Ultraculture complete medium for transfection of primary neurons). 24 h later the first batch of supernatant was collected, cells were overlaid with 9 ml D10 medium (or Ultraculture complete medium for transfection of primary neurons), and after another 24 h the second batch of supernatant was collected and combined with the first batch. Lentiviral supernatant was frozen in aliquots and stored at -80°C. Prior to use, lentiviral supernatant was centifuged for 5 min at 2,000 g.

#### Ultraculture complete medium (50 ml)

48.25 ml Ultraculture serum-free medium (Lonza BP12-725F), 0.5 ml 100 mM sodium pyruvate (Lonza 13-115E), 1 ml penicillin/streptomycin (Gibco 15070-63, 5,000 u/ml), 0.25 ml GlutaMAX (Thermo Fisher 35050061).

#### D10 medium (500 ml)

450 ml DMEM (Gibco 11965-092), 10% heat-inactivated FBS (GE Life Sciences SH30396.03), 10 ml penicillin/streptomycin (Gibco 15070-63, 5,000 u/ml), 5 ml L-glutamine (Gibco 25030-081).

### Mouse cortical neuron culture and transduction

Cas9^+/+^ mouse cortical neurons were cultured following published protocols, with modifications ^53,54^.

#### Brain dissociation

Brains from E15 or E16 Cas9^+/+^ embryos were collected in 15 ml conical tubes filled with L-15 medium (Thermo Scientific 11415064). Brains were allowed to settle at the tube bottom by gravity, supernatant was removed, and the tube was again filled with L-15. The washing procedure was repeated for a total of three times. After the last wash, all liquid was removed, and TrypLE (Thermo 12605010; containing recombinant trypsin and EDTA) was added (1 ml per brain). The tube was gently inverted and immediately submerged in a 37°C water bath for 5-6 min. In our hands, limiting trypsin digestion to no more than 6 minutes was critical to obtain healthy neuron cultures. During the incubation period, the tube was inverted briefly for 2 or 3 times. After incubation at 37°C, DNase I (25 µl per brain, stock 10 mg/ml) (Sigma 10104159001) was added, the tube inverted, and trypsin action was stopped by adding 1 volume of plating medium (see below) to 1 volume of brain suspension. Brains were pelleted by centrifugation (200 g, 1 min) and washed two times with L-15 (centrifugation at 200 g, 1 min). The use of serological pipettes must be avoided at this stage as brains tend to strongly stick to polystyrene. After the final wash, all supernatant was removed, and 1 ml plating medium was added.

#### Trituration

Trituration was carried out using a standard 1 ml polypropylene pipet tip, pipetting the cell suspension up and down with varying degrees of shearing. The first 3 passes (1 pass = pipetting up and down) were carried out without shearing, followed by 7-10 passes with shearing. The shearing force was varied depending on the cloudiness of the solution. After trituration, 9 ml plating medium was added, the cell suspension was centrifuged at 200 g for 5 min, cells were resuspended in plating medium (1 ml per brain), and passed through a 40 um cell strainer. Cells were counted by trypan blue exclusion (cell yield per brain is approximately 10 x 10^6^ cells).

#### Plating

Cells were plated in poly-D-lysine-coated culture dishes (see below) at a density of 180,000 to 200,000 cells per cm^2^ in plating medium (e.g. 0.6 ml per well of a 24-well plate, with a well area of 1.9 cm^2^). After cell adhesion for 1-2 h in a 37°C, 5% CO2 incubator, the plating medium was aspirated and immediately replaced by Neurobasal Plus complete medium (see below) (0.6 ml per well of a 24-well plate). Aspiration and media addition must be carried out rapidly, as cells tend to dry out quickly.

#### Lentiviral transduction

were carried out on DIV 1 by adding lentiviral supernatant (collected in Ultraculture complete medium, see below) to neuronal cultures. Final supernatant concentration was 20% (vol/vol) or lower.To prevent proliferation of nonneuronal cells, mitotic inhibitors FUDR (Sigma F0503) and uridine (Sigma U3003) were added on DIV 1 to a final concentration of 6-8 μM each (stock was 10 mM FUDR and 10 mM uridine in H_2_O). Every 2-3 days, about 1/3 to 1/2 of culture media was replaced with fresh Neurobasal Plus complete medium, with no further addition of mitotic inhibitors. Media evaporation was minimized by filling spaces between wells with H_2_O, and by keeping culture dishes on top of water-filled trays inside the incubator.

#### Coating of culture dishes

Poly-D-lysine (0.1 mg/ml, Thermo Fisher A3890401) was dispensed into wells (e.g. 0.6 ml per well of a 24-well plate), and plates were incubated at 37°C for 1 h to 24 h. After incubation, wells were washed 4 to 6 times with sterile, nuclease-free water. After the final wash, all liquid was aspirated thoroughly and plates were air dried. Plates can be used immediately or wrapped and stored at 4°C for up to 7 days.

#### Plating medium (50 ml)

45.583 ml MEM without L-glutamine (Thermo Fisher 11090081), 2.5 ml (5% final) heat-inactivated FBS, 0.667 ml (0.6% or 33.33 mM final) 45% D-glucose (Sigma G8769), 0.25 ml 200 mM GlutaMAX (Thermo Fisher 35050061), and 1 ml penicillin/streptomycin (final 100 u/ml penicillin, 100 ug/ml streptomycin; Gibco 15070-63).

#### Neurobasal Plus complete medium (50 ml)

47.75 ml Neurobasal Plus medium (Thermo Fisher A3653401), 1 ml B-27 Plus supplement (Thermo Fisher A3582801), 1 ml penicillin/streptomycin (final 100 u/ml penicillin, 100 ug/ml streptomycin; Gibco 15070-63), 0.25 ml GlutaMAX(200 mM, Thermo Fisher 35050061).

### Dorsal root ganglion neuron culture and transduction

Cultures of mouse primary sensory neurons were prepared using dissociated DRGs from E13 or E14 mouse embryos, according to published protocols ^52^. Cas9^+/+^ embryos were used for experiments involving sgRNAs and for some overexpession experiments; for some of the latter, wild-type embryos were used.

### Dissociation

DRGs from up to 8 E13 or E14 mouse embryos were collected in a sterile 2 ml tube in L-15 media (Thermo Scientific 11415064) and washed three times in L-15 (centrifugation at 200 g for 30 seconds). After removing the supernatant completely, TrypLE (Thermo 12605010; containing recombinant trypsin and EDTA) was added (1 ml for DRGs from up to 8 embryos), and the tube was immediately submerged in a 37°C water bath and incubated for 5-6 min (during incubation, the tube was briefly inverted 2 or 3 times; keeping digestion time below 6 min was found to be essential). Immediately after digestion, 800 µl complete DRG medium (see below) was added, and DRGs were pelleted at 300 g for 30 seconds. DRGs were then washed 3 times in complete DRG medium (centrifugation at 300 g for 30 seconds). After the last centrifugation, supernatant was completely removed, and 1 ml complete DRG medium was added.

### Trituration

A 1 ml standard plastic tip was used for trituration, coated with 10% FBS in complete DRG medium to keep DRGs from sticking. DRGs were first resuspended with 3 pipetting passes (1 pass = up + down), followed by up to 20 passes with medium to harsh shearing. Cell suspension cloudiness served as a guide for the number of passes and shearing force applied. Cells were counted immediately after trituration (cell counts were in the range of 10^6^ cells per 6-8 embryos, with broad variability). After another pelleting step at 300 g for 3 min, cells were resuspended in complete DRG medium at a concentration of 10,000 cells/ul.

### Spot plating

When cells were ready for plating, laminin solution was aspirated from prepared culture plates (24-well or 48-well plates), and after a brief wash with sterile H_2_O all liquid was completely aspirated from culture wells. Working rapidly (laminin is thought to be sensitive to drying), dissociated DRG neurons were spot plated by dispensing 2.5 µl cell suspension containing 25,000 cells total in the well center (evaporation was minimized by filling spaces between wells with sterile H_2_O, and during cell dispension the culture dish was set atop a large tray of water). Plates were immediately transferred to a 37°C, 5% CO_2_ humidified incubator, and cells were allowed to adhere for 6 min. Wells were gently filled with complete DRG medium (24-well: 0.6 ml/well; 48-well: 0.3 ml/well), and plates were transferred back to the incubator. <u>Sparse plating.</u> For some experiments, DRG neurons were sparsely plated by diluting cells in complete DRG medium, aiming at a plating density of 25,000 cells/cm^2^ (in our experience, DRG neurons looked unhealthy at plating densities of 50,000 cells/cm^2^ and higher).

### Culture plates

24-well or 48-well plates were used for DRG neuron culture. Wells were coated sequentially with poly-D-Lysine (PDL, Fisher Scientific A3890401) and laminin (Thermo Scientific 23017015). First, PDL was dispensed into each well (24-well: 350 µl; 48-well: 200 µl), and plates were kept for 1-24 hours at 37°C. Plates were then washed with PBS for 4-6 times (washing with PBS instead of H_2_O provides better pH stability for PDL), and laminin solution (5 ug/ml diluted in PBS w/o Ca^++^ w/o Mg^++^) was added to wells (24-well: 350 µl; 48-well: 200 µl). Plates were incubated at 37°C for 3-5 hours. Laminin was only removed prior to cell plating (see below).

### Complete DRG medium (50 ml)

47.25 ml Neurobasal medium (Thermo Scientific 21103049), 1.0 ml B-27 supplement 50x (Gibco 17504044), 0.25 ml GlutaMAX 200 mM (Thermo Fisher 35050061), 1 ml penicillin/streptomycin (final 100 u/ml penicillin, 100 ug/ml streptomycin; Gibco 15070-63), 0.5 ml 2.5 M glucose (final 25 mM; Sigma G8769), 25 µl 100 ug/ml NGF 2.5S (final 50 ng/ml; Promega G5141).

### Lentiviral transduction

were carried out as described for cortical neurons, with the obvious difference of using complete DRG medium.

### Axotomy and cell imaging

Neurites of cultured mouse DRG neurons were axotomized on DIV 11-13, using a 2.75 mm blade mounted to a handle (Fine Science Tools 10035-10 and 10035-12), or larger off the shelf hobby knife blades. Axotomy was performed on the stage of an inverteded fluorescence microscope, on neurites marked by EGFP or mCherry cytoplasmic markers. Observing cells under the microscope while cutting gave excellent control over knife positioning, avoiding stray cell bodies, and completeness of introduced cuts. Typically, half of neurites in a well were severed from their cell bodies (‘cut’ condition), while the other half was left intact (‘uncut’ condition).

For time lapse imaging of neurodegeneration in live neurons, we used the Zeiss Celldiscoverer 7 automated microscope. Its features include temperature and CO_2_ control, software and hardware-based focus, LED-based fluorescence imaging capabilities, and a large array of programmable settings for acquisition. For axotomy-induced neurodegeneration, conditions were set up in triplicate wells (or duplicates in rare instances). For each well, 3 or 4 images of uncut, and 3 or 4 images of cut neurites were imaged at multiple time points (typically ranging from 4 h to 24 h post axotomy).

### Alternative Sarm1 activation methods

#### B/B homodimerizer

(AP20187) (Medchemexpress HY-13992) was dissolved at 1 mM in anhydrous DMSO and stored at - 20°C.

#### Vacor (Pyrinuron)

(Chem Service N-13738-100MG) was dissolved at 10 mM in 0.04 M HCl and stored at -80°C.

### HIF-1 alpha subunit immunofluorescence staining

Mouse cortical neurons from Cas9^+/+^ embryos were plated at 200,000 cells/cm^2^ in 96-well black plates (µ-Plate 96 Well Black, ibidi #89626)(see above for culture method), and transduced on DIV 1 with indicated guides or constructs. Cells were fixed and stained on DIV 13.

#### Fixation

All media was completely removed, 100 µl of 4% PFA in PBS was added per well, and cells were incubated at room temperature for 15 min. The fixative was removed, and 100 µl of 0.2% TX-100 in PBS was added per well and left for exactly 1 min. After removing the TX-100 solution, cells were washed once with 100 µl PBS.

#### Primary antibody

Anti-HIF-1 alpha chain antibody (Cell Signaling Technology #96169T, HIF-1alpha (D1S7W) XP rabbit mAb) was diluted 1:400 in 0.02% TX-100 in PBS. Fixed cells were incubated with 80 µl of antibody solution for 2 hours, followed by two washes with 100 µl of 0.02% TX-100 in PBS.

#### Secondary antibody

Donkey anti-rabbit Cy3 conjugate (Jackson 711-165-152) was diluted 1:250 in 0.02% TX-100 in PBS. Cells were incubated with 100 µl secondary antibody solution for 1 hour, then washed twice with 100 µl of 0.02% TX-100 in PBS.

Cells were overlaid with 0.4 ug/ml Hoechst 33342 in 0.02% TX-100 in PBS, incubated for 30 min, and imaged on a Zeiss Axiovert Observer upright fluorescence microscope.

### Sarm1 Western blot

Mouse cortical neurons were plated at 190,000 cells/cm^2^ in 6 cm dishes (see above for culture method) and transduced on DIV 1 with indicated lentiviral constructs. Lysates were prepared on DIV 11. Culture media were aspirated and cells were washed briefly with PBS (2 ml per 6 cm dish). After complete aspiration of PBS, lysis buffer (150 µl per 6 cm dish, see below) was added to cells, making sure all cells were covered. Working in the cold room, plates were held at an angle, and the lysed cell lawn was pushed to one side of the dish using a cell lifter. The lysate was briefly homogenized by pipetting up and down, then transferred to a 1.5 ml tube. Working at room temperature, each lysate of ∼150 µl was mixed with 50 µl of 4x Laemmli buffer (see below), briefly vortexed, mixed with 0.5 µl benzonase (Sigma #E8263-5KU) and vortexed again, then kept at room temperature for 5 min. Lysates were heated to 95°C for 5 min, spun briefly, and frozen at -20°C.

Proteins were electrophoretically separated by SDS-PAGE, loading 10 µl lysate per slot (corresponding to ∼10,000 originally plated cells) of a 4-20% Tris-Glycine Mini Protein Gel (Life Technologies/Invitrogen #EC60285BOX). Proteins were transferred onto 0.45 um nitrocellulose by semidry blotting and visualized with Ponceau S (1% Ponceau S, 1% glacial acetic acid in H_2_O). Sarm1 was detected using rabbit monoclonal anti-Sarm1 antibody (Cell Signaling #13022S) diluted 1:800 in TBS-T with incubation for 3 hours at room temperature, followed by anti-rabbit HRP conjugate (Jackson #711-035-152) diluted 1:2000 in TBS-T for 1 hour. ECL solution (Super Signal West Dura, Thermo #34075) and autoradiography film were used to detect the signal.

#### Lysis buffer

50 mM Tris HCl pH 7.5, 150 mM NaCl, 0.1% SDS, 1% NP-40, 1% DDM, Complete protease inhibitor (Roche #11697498001), 1 mM DTT

#### 4x Laemmli buffer

200 mM Tris HCl pH 6.8, 8% SDS, 40% glycerol, 400 mM DTT, 0.02% bromophenol blue

#### Electrophoresis buffer

25 mM Tris pH 8.3, 192 mM glycine, and 0.1% SDS Transfer buffer: 25 mM Tris, 192 mM NaCl, 15% methanol

#### TBS-T

20 mM Tris pH 7.4, 150 mM NaCl, 0.1% Tween-20

### Neurite integrity index (NII)

A detailed description of our custom neurite integrity index (NII) computation method, along with code repository, sample images, and step-by-step instructions for image processing in Python and R, can be found at https://github.com/pmeran/neuriteX/

### Genome-wide sgRNA screens

#### Preparation of screening libraries in HeLa and HEK293T cells

The plasmid pool for the genome-wide human sgRNA library “Brunello” (Addgene #73178) ^18^ comprises a total of 76,441 guides, plus 1000 non-targeting control guides. Each of the 19,114 targeted genes is covered by 4 non-overlapping guides. The library comes subcloned into vector lentiGuide-Puro (Addgene plasmid #52963) which is puromycin-selectable. The plasmid library was amplified following a protocol optimized to preserve guide complexity (https://www.addgene.org/protocols/pooled-library-amplification/). After packaging the library into lentivirus (see above), multiple aliquots of virus supernatant were frozen at -80°C. The titer was determined for both HeLa and HEK293T cells as targets.

SARM1ΔARM is a truncated version of SARM1 lacking the autoinhibitory ARM domain ^57^, thereby exhibiting constitutive NAD^+^ hydrolase activity and mimicking the effects of activated full-length SARM1 in neurons ^17,58^. Expression of dSarmΔARM in fly neurons in vivo ^15^ causes neurite degeneration and ultimately cell death. Similarly, expression of SARM1ΔARM in mammalian cell lines leads to cell death. We hypothesized that genes required for SARM1ΔARM-induced cell death could be identified in genome-wide CRISPR-Cas9 sgRNA screens.

The screens were conceived as a genome-wide survival screens. To generate screening libraries, we first prepared Cas9-transduced HeLa and HEK293T cell lines. In a second step, we transduced cells with doxycycline-inducible SARM1ΔARM (using in the multicistronic construct SARM1ΔARM-T2A-EGFP-P2A-BSD, with EGFP as a reporter, and BSD (blasticidin S deaminase) for optional chemical selection of integrated SARM1ΔARM), expanded clonal populations thereof, and verified that the sequence of chromosomally integrated SARM1ΔARM was correct. Cloning and sequence verification of integrated SARM1ΔARM were necessary because lentiviral genomes suffer from high error frequencies. Earlier library versions relied on transduction of SARM1ΔARM from a lentiviral pool without deriving clones. This approached turned out to be highly flawed, because due to high error rates in lentiviral genomes, a sizeable fraction of integrated SARM1ΔARM sequences carried attenuating mutations (many of them identical or similar to the commonly used E642A NADase-dead mutant) and did not kill cells when induced with doxycycline, leading to rampant accumulation of false-positive sgRNAs in the cell population surviving the screen.

Third, cell lines were transduced with the human genome-wide sgRNA library “Brunello” in lentiviral vector lentiGuide-Puro (Addgene #73178) ^18^. This sgRNA library targets the human genome with 4 guides per gene and comprises about 78,000 guides total. Library transduction was performed at a MOI of 0.25 or lower so that the overwhelming majority of transduced cells carried only one sgRNA. Nontransduced cells were eliminated efficiently with puromycin, the selection marker on lentiGuide-Puro.

Gene editing with CRISPR-Cas9 in cell lines typically reaches a plateau 12 to 14 days after guide introduction. SARM1ΔARM was therefore induced with doxycycline 2 weeks after sgRNA library transduction, in a library population of 80 x 10^6^ cells (overrepresenting roughly 80,000 guides by a factor of 1000). Surviving clones were expanded for 2-3 weeks to obtain sizeable populations for genomic DNA preparation and guide repertoire analysis.

### Bioinfomatic analysis of CRISPR-Cas9 screens

#### Sample preparation

Guide repertoires of pre-selection libraries and expanded post-selection populations were analyzed using the Illumina MiSeq platform, following published protocols ^59^, with modifications. Genomic DNA of pre-selection libraries in HeLa and HEK293T was prepared for 100 x 10^6^ cells of each cell library using the Blood & Cell Culture DNA Maxi Kit (Qiagen 13362). Genomic DNA for populations surviving screen selection (typically amplified to millions of cells) was prepared with the DNeasy Blood & Tissue Kit (Qiagen 69506). Genomically integrated sgRNA cassettes were amplified by PCR in two steps, yielding amplicons with a total length of 373 bp (with a maximum stagger length of 8 bp). PCR primers for PCR 1 and PCR 2, and the full amplicon sequence with annotations are listed below. PCR 2 introduces index combinations that allow for multiplexing of multiple samples in one run. PCR amplicons were column-purified, quantitated by Qubit, mixed at ratios corresponding to estimated guide repertoire complexities, and loaded onto the MiSeq flow cell (Illumina MiSeq Reagent Kit v2, MS-102-3001).

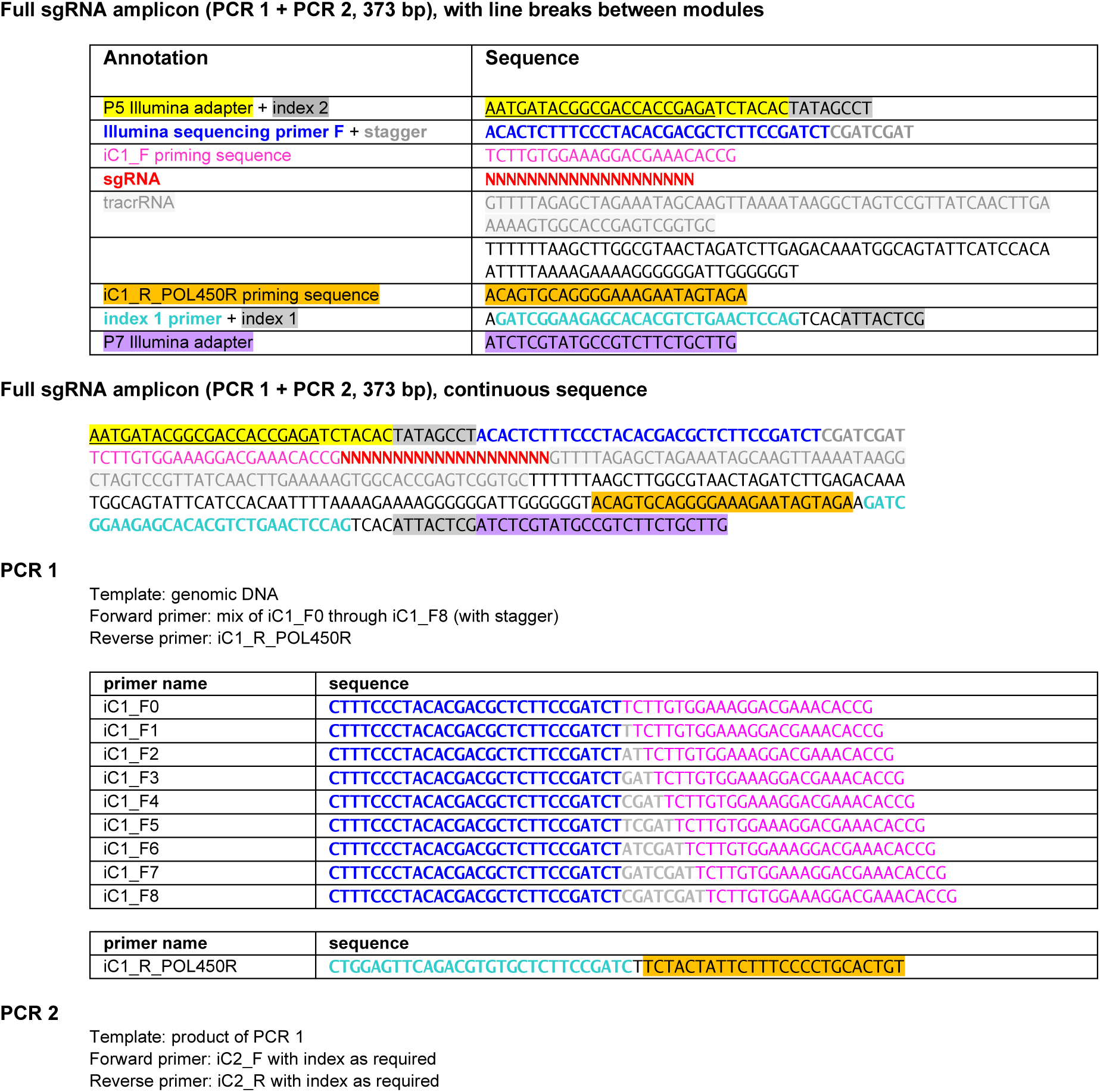
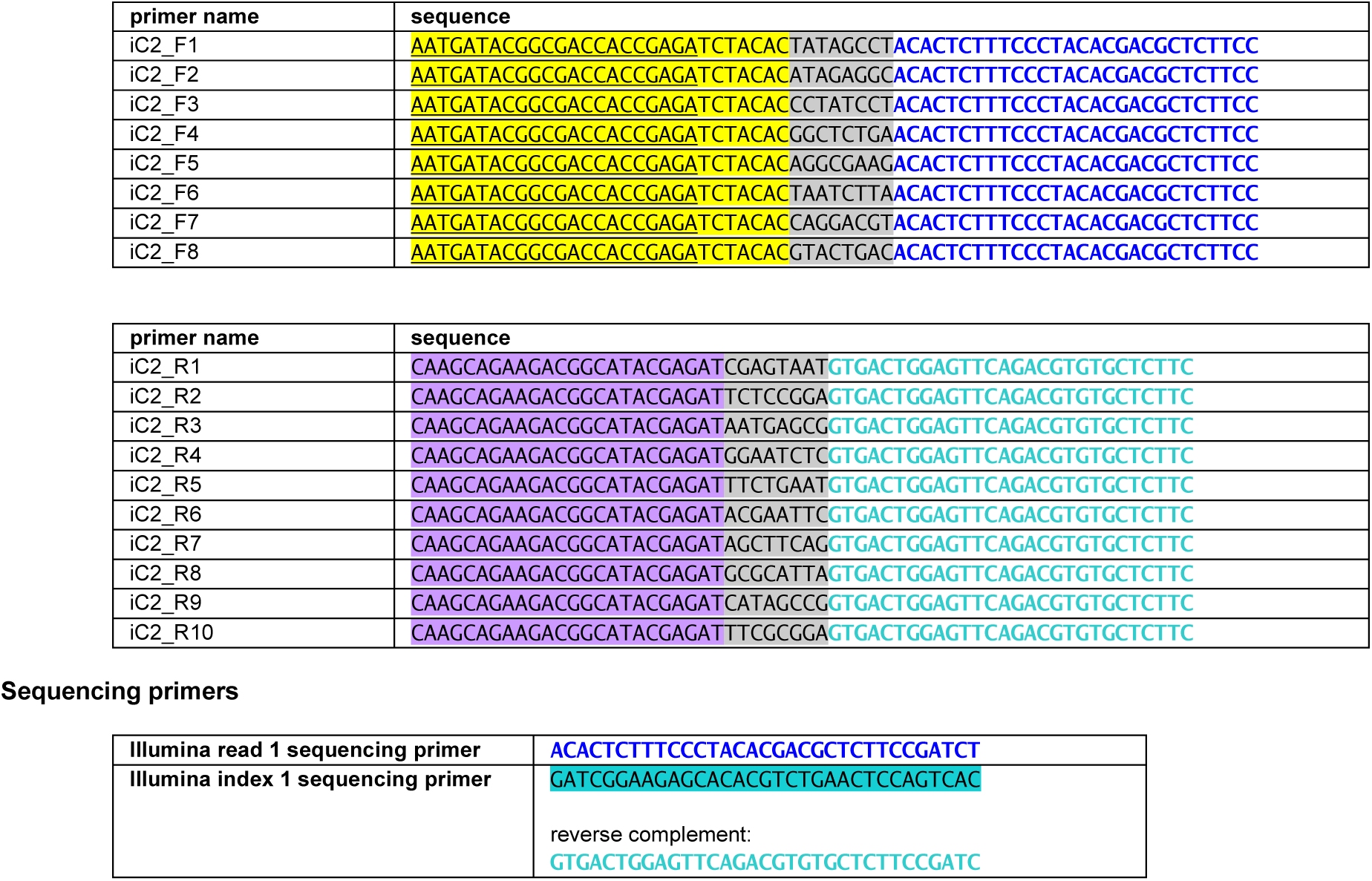

#### Analysis of sequencing data

Raw reads in fastq format were transferred to the OHSU ACC Exacloud compute cluster with a Linux command line environment that allows convenient handling of large datasets. Guide sequences were extracted from fastq reads based on sequence context and demultiplexed using Linux bash scripts, then aligned to the human genome with bowtie2 ^60^. The resulting alignment files in .sam format were processed with the samtools command line module, and lists of read counts for all retrieved guides were generated. PBNPA (<u>p</u>ermutation-<u>b</u>ased <u>n</u>on-<u>p</u>arametric <u>a</u>nalysis) ^61^ is a tool for CRISPR screen analysis that runs on the popular R statistics platform. We chose PBNPA because it performs well with screen data containing single replicates, a screen design we deliberately chose to fit a larger number of experimental conditions into the available read space. PBNPA transforms read counts for guides into gene-level metrics, computing p values as a measure of statistical significance. For convenient exploratory analysis, we created interactive charts with the Python module plotly, plotting the statistical significance of a gene (expressed as -log10(p)) versus its enrichment (expressed as log2(mean fold change)). Static charts for selected gene sets were created in R with ggplot2.

### Sorting of primary mouse neuron nuclei and RNA-Seq

Cultures of mouse cortical neurons and DRG neurons were routinely supplemented with mitotic inhibitors (FUDR and uridine 6-8 μM each on DIV 1 only) to suppress glial overgrowth. However, a large population of non-dividing non-neuronal cells persisted, as evident from staining with nuclear dye Hoechst 33342. To obtain RNA-Seq profiles of a clean neuronal population, we isolated nuclei by flow cytometry as the starting material for RNA-Seq, rather than bulk lysates. To identify nuclei from neurons with high specificity, cells were transduced with a fluorescent reporter fused to the nuclear protein Lamin-B1, driven by the human synapsin 1 promoter (P.Syn::EGFP-Lamin-B1).

Cultures of mouse cortical neurons were started at a plating density of about 150,000 cells/cm^2^. DRG neuron cultures were sparsely plated at ∼25,000 cells/cm^2^ (plating densities of 50,000/cm^2^ and higher led to clear signs of degradation over time). Nuclei were isolated following established protocols ^62^. In brief, after a total culture period of 12 days, all media were aspirated, followed by immediated addition of ice-cold buffer H (= homogenization buffer: 10 mM Tris pH 8.0, 250 mM sucrose, 25 mM KCl, 5 mM MgCl_2_, 0.1% Triton X-100, 0.5% RNasin Plus RNase inhibitor (Promega), 1x EDTA-free Complete potease inhibitor (Roche), 0.1 mM DTT), about 800 µl per 10 cm dish. Cells were incubated at 4°C in buffer H for 10-15 min, scraped off with a cell lifter, and sheared gently by pipetting up and down 5-10 times. After centrifugation to pellet nuclei (700 g, 8 min, 4°C), the supernatant was discarded and nuclei were resuspended in 1 ml of ice-cold buffer PBS-P (PBS with 0.8% nuclease-free BSA, 0.5% RNasin Plus RNase inhibitor, and 0.5 ug/ml Hoechst 33342). Nuclei were kept on ice until sorting. Nuclei were morphologically stable for several hours as of microscopic inspection, and post-sort nuclear RNA was intact based on Bioanalyzer profiling.

For flow cytometric sorting, the gate was drawn around Hoechst/EGFP high positive events. Using a 100 um nozzle, nuclei were sorted directly into PBS-P (with RNasin Plus RNase inhibitor reduced to 0.1%). Per sample, about 500,000 nuclei were collected for mouse cortical neurons, compared to 15,000 nuclei for the less abundant DRG neurons. Nuclei were subsequently pelleted by centrifugation (800 g, 10 min, 4°C), and nuclear total RNA was isolated using commercial kits (Qiagen RNeasy Plus mini kit for mouse cortical neurons; Zymo Quick-RNA Microprep kit for DRG neurons). For both cell types, the obtained total RNA was of sufficient quantity (above 25 ng total RNA for sequencing library preparation without additional amplification step), and of sufficient quality based on Bioanalyzer profiles (the RIN = RNA integrity number was close to 1 for cortical neuron samples, and over 0.9 for most DRG neuron samples; a RIN > 8.5 is generally considered satisfactory). Our sequencing core used the Takara Smart-Seq v4 Plus kit for library preparation, and the Illumina NovaSeq 6000 sequencer to collect 50 million read pairs per sample.

For both mouse cortical neurons and DRG neurons, two conditions were compared: cells transduced with empty vector backbone, and with Hif1a-S. For each condition, 3 biological replicates were set up and analyzed.

### RNA-Seq data analysis

RNA-Seq data analysis was performed on the OHSU exacloud compute cluster in a Linux environment, and on a commercial PC using the R statistics package and custom scripts in Python and Javascript. The following software modules were used along the analysis pipeline. Mouse genome and transcriptome data (GRCm39) were downloaded from www.ensembl.org. Original .fastq files generated by the Illumina sequencer all passed FastQC quality control, and were aligned to mouse genome and transcriptome using the STAR aligner (version 2.7.11b) ^63^ to create .bam alignment files. The transcript quantification tool Salmon (v1.10.0 ^64^) served to convert .bam alignment files to transcript-level count matrices. Switching to R, Salmon-generated count matrices were imported with tximport (version 1.30.0) into the DESeq2 differential gene expression analysis tool (version 1.42.0, ^65^). Basic RNA-Seq quality checks with DESeq2 (principal component analysis, dispersion estimation, total sample counts, library complexity, number of detected genes) suggested that the data were of high quality.

Using Javascript, a custom tool was developed for interactive data exploration in a web browser. We plan to share the respective website upon request in a future peer-reviewed version of this article. Our analysis tool allows to interactively explore the full RNA-Seq datasets for mouse cortical neurons and DRG neurons. Gene symbols are used as main gene reference, while ensembl gene IDs are shown where available. The shown numerical values – TMPs, log2-fold change, and p values – were all generated by Salmon and DESeq2. Alongside RNA-Seq data, the tool shows a large collection of annotated gene sets that is based on the Molecular Signatures Database for mouse (MSigDB version 2023.2, freely available from www.gsea-msigdb.org). MSigDB includes gene sets curated by different organizations and platforms, compiled based on morphological, functional, and biochemical criteria. Gene sets can be searched, selected, and used to filter RNA-Seq data, with output displayed in graphical and tabulary form.

Raw RNA-Seq .fastq files are available on the NIH Sequence Read Archive (https://www.ncbi.nlm.nih.gov/sra) under accession number PRJNA1197120. Processed RNA-Seq data in excel spreadsheet format are provided in Supplementary File “RNA-Seq”. A web-based tool under https://d3259ptldlzjvg.cloudfront.net/ (currently password-protected; access credentials can be provided upon request) allows comparative exploration of processed RNA-Seq data for the two investigated neuron types.

### Quantitation of NAD+ and NADH in mouse neurons

NAD+ and NADH concentrations were measured in cells using a bioluminescence assay (NAD/NADH-Glo Assay, Promega #G9071), following the manufacturer’s instructions. Mouse cortical neurons were plated in 96-well plates (µ-Plate 96 Well Black, ibidi #89626; well area 0.56 cm^2^) on DIV 0 at a density of 190,000 cells/cm^2^, and transduced with empty vector or Hif1a-S on DIV 1. On DIV 12, Vacor was added at indicated concentrations in a staggered fashion, at time points -6 h, -3 h, and -1 h, and cells were lysed simultaneously at t = 0. One of the plates was used for imaging and neurite integrity index (NII) computation.

Cells were processed as follows. Immediately after complete media removal, 30 µl PBS and 30 µl 1% DTAB in base solution (0.2 M NaOH) was added to each well, and lysates from two identical wells were pooled (bringing the number of replicates from 6 to 3, and the volume in each well to 120 ul). Lysates were then split into two identical portions of 50 µl each and transferred to a transparent 96-well polystyrene plate. Half of that plate was used for measurement of NAD+, and the other half – with identical samples and layout – for NADH. To wells slated for NAD+ measurement, 25 µl of 0.4 M HCl was added. The entire plate was incubated at 60°C for 15 min and cooled to room temperature. Wells for NAD+ measurement received 25 µl of 0.5 M Tris to neutralize the pH; while wells for NADH measurement received 50 µl of a 1:1 mixture of 0.4 M HCl and 0.5 M Tris. At this stage, the volume in each well was 100 µl. For measurement, 50 µl from each well was transferred to an opaque, white 96-well plate, and mixed with 50 µl of NAD/NADH-Glo detection reagent. After a 30 min incubation, luminescence was quantified using the ClarioStar Plus microplate reader (BMG Labtech). Each condition was quantified in triplicates, and recorded counts were normalized using time point 0 as reference.

### Generation of genetically modified flies

The MARCM genetic modification system (Mosaic Analysis with a Repressible Cell Marker ^39,40^) allows the targeted expression of virtually any gene in fly cells defined by a specific promoter, including sparse labeling and introduction of homozygosity in the targeted cells. Expression of mouse Hif1a-S in fly wing sensory neurons was achieved using the Gal4/UAS transcription factor/promoter system in the sensory neuron-specific OK371-Gal4 driver line (which the glutamatergic neuron-specific OK371 promoter), with UAS-tdTomato added as reporter ^66^. Two fly strains were generated, one with UAS-Hif1a-S on the 2^nd^ chromosome (site attp40) and the other with UAS-Hif1a-S on the 3^rd^ chromosome (site attp154), to provide greater flexibility for incorporating additional genetic features through crossing.

#### Generation of *5xUAS-mHif1a-S* flies

The open reading frame of the undegradable mouse Hif1a (Hif1a-S) was PCR-amplified while incorporating *XhoI* and *NheI* sites at the 5’ and 3’ ends, respectively. The amplified fragment was then cloned into the *pattB-5xUAS* vector, the sequence was confirmed and the DNA was injected into *y*^1^ *w*^67c23^ *P{CaryP}attP18* (RRID:BDSC_32107) and *y1 w*^67c23^*; P{CaryP}attP154* flies by BestGene.

Fly lines used in this study:

|  |  |  |
| --- | --- | --- |
| *OK371-gal4* | Bloomington Drosophila Stock Center (BDSC) | RRID:BDSC_26160 |
| *10XUAS-ivs-myrTdTomato* | BDSC | RRID:BDSC_32222 |
| *5XUAS-LacZ* | BDSC | RRID:BDSC_3956 |
| *aseFLP<sup>2c</sup>* | Freeman Lab (Neukomm et al., 2014) | N/A |
| *aseFLP<sup>3b</sup>* | Freeman Lab (Neukomm et al., 2014) | N/A |
| *tub-gal80 FRT19A* | BDSC | RRID:BDSC_5132 |
| *FRT19A* | BDSC | RRID:BDSC_1709 |

## Author contributions

Paul Meraner performed experiments, analyzed CRISPR-Cas9 screens and RNA-Seq data, acquired and analyzed axotomy imaging data, wrote image analysis software, instructed and supervised technicians, contributed to concepts and directions, and co-wrote the manuscript.

Adel Avetisyan performed fly experiments, made general contributions to experiment design and interpretation, and co-wrote the manuscript.

<u>Kevin Swift</u> and <u>Ya-Chen Cheng</u> provided technical help and performed experiments in the areas of molecular biology and cell culture. Ya-Chen Cheng helped with mouse husbandry.

Romina Barria provided technical support in experiments involving molecular biology and cell culture, and helped with mouse husbandry.

Marc Freeman initiated the study, provided institutional and financial support, set overall goals and directions, supervised collaborations, coordinated experimental efforts, and co-wrote the manuscript.

All authors read and consented to the final version of the manuscript.

## Glossary

**HIF-1**, hypoxia-inducible factor 1. HIF-1 is a heterodimeric transcription factor complex composed of alpha (gene *Hif1a*) and beta subunit (gene *Arnt*). A variant of *Hif1a* with 3 mutations (P402A/P577A/N813A based on mouse transcript NM_010431.3) escapes constitutive ubiquitination and proteasomal degradation, and is here referred to as Hif1a-S (S for “stabilized”). Similarly, a HIF-1 complex incorporating Hif1a-S alpha subunit (and a wild-type beta subunit) is referred to as HIF-1-S. The terms “stabilized HIF-1”, “HIF-1-S”, and “Hif1a-S” are used interchangeably. **DIV**, day in vitro. **NII**, neurite integrity index (1.0 for intact neurites, 0.0 for degenerated neurites). **DRG**, dorsal root ganglion. **TPM**, transcripts per million. **sgRNA**, single guide RNA (sometimes referred to simply as “guide”). **dpe**, days post eclosion. **dpa**, days post axotomy. Genes are *italicized* to emphasize reference to the the gene. Following common practice, human genes are given in capital letters, and mouse genes with only the first letter capitalized.

## Supporting information

Sarm1 Supplementary Figures & Tables.pdf

Sarm1 Supplementary File gene_sets.xlsx

Sarm1 Supplementary File RNA-Seq.xlsx

## Acknowledgments

We express our gratitude to the following colleagues and institutions: Yibing Jia for operating the Illumina MiSeq sequencer at the ONPRC Molecular Biology Core at the OHSU West Campus; the OHSU Integrated Genomics Laboratory (IGL); the OHSU flow cytometry core; the OHSU Advanced Computing Center (ACC) for supporting use of the OHSU Exacloud compute cluster; Stefanie Kaech-Petrie from the OHSU Advanced Light Microscopy Core for help and support with the Zeiss Celldiscoverer 7 automated microscope.

For insightful discussions our thanks go to Julie Soutourina (CNRS, Université Paris Saclay, Gif-Sur-Yvette, France), Dylan Taatjes (University of Colorado, Boulder, CO), Michael Coleman (University of Cambridge, Cambridge, UK), and Juan Bolaños (University of Salamanca, Salamanca, Spain).

This work was supported by NIH/NINDS grants RO1 NS059991 (to MRF), a sponsored research agreement from Nura Bio, OHSU and the HHMI.

