## Supplementary material for "Hypoxia-inducible factor 1 protects neurons from Sarm1-mediated neurodegeneration": Sarm1 Supplementary Figures & Tables.pdf

### Supplementary Fig. S1

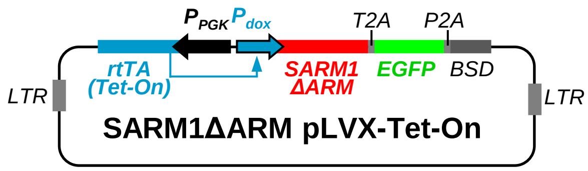


**Supplementary Fig. S1 | Lentiviral SARM1ΔARM construct used in CRISPR-Cas9 survival screens**
Promoter $P_{PGK}$ drives constitutive expression of the reverse tetracycline-controlled transactivator (rtTA), which must bind doxycycline to drive expression from the doxcycycline-inducible promoter ($P_{dox}$). Coding sequences for SARM1ΔARM, EGFP, and blasticidin resistance (BSD = blasticidin S deaminase) are separated by autocatalytic T2A and P2A peptides.

### Supplementary Fig. S2

**a**

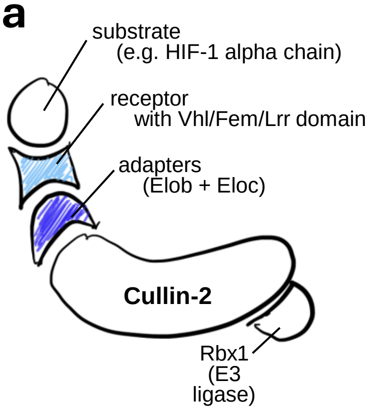


**b**

| | gene family | subfamily | gene | protein |
|---|---|---|---|---|
| scaffold | | | **Cul2** | Cullin 2 |
| adapters | | | **Elob** | Elongin-B |
| | | | **Eloc** | Elongin-C |
| receptors | Vhl | | **Vhl** | von Hippel-Lindau disease tumor suppressor |
| | Fem | | **Fem1a** | |
| | | | **Fem1b** | |
| | | | **Fem1c** | |
| | Lrr | Lrr | **Lrr1** | |
| | | Lrriq (3 genes) | **Lrriq1** | Leucine-rich repeat and IQ domain-containing protein 1 |
| | | | **Lrriq3** | Leucine-rich repeat and IQ domain-containing protein 3 |
| | | | **Lrriq4** | |
| | | Lrrtm (4 genes) | **Lrrtm1** | |
| | | | **Lrrtm2** | |
| | | | **Lrrtm3** | |
| | | | **Lrrtm4** | |
| | | Lrrtc (~ 60 genes) | | |

**Supplementary Fig. S2 | Composition of Cullin-2 RING E3 ubiquitin ligases (CRL2)**
**a** Schematic of the Cullin-2 RING E3 ubiquitin ligase complex.
**b** List of CRL2 adapters and substrate receptors.

### Supplementary Fig. S3

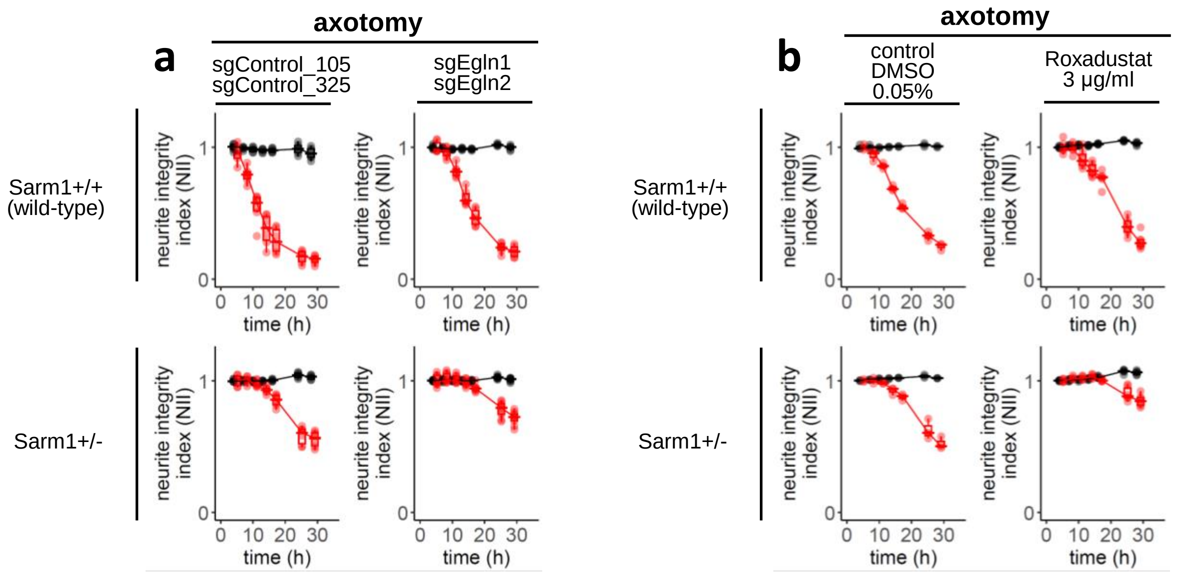


**Supplementary Fig. S3 | Suppression of prolyl hydroxylases moderately protects neurons from Sarm1-mediated neurodegeneration**

**a Cas9-mediated ablation of prolyl hydroxylases Egln1 and Egln2.** DRG neurons from Sarm1+/+ mice (top panels) or from Sarm1+/- mice were transduced on DIV 1 with control guides (sgControl_105, sgControl_325) or guides targeting prolyl hydroxylases Egln1 (sgEgln1) and Egln2 (sgEgln2). Axotomy was performed on DIV 11. Charts show the neurite integrity index (NII) plotted against time.

**b Pharmacological inhibition of prolyl hydroxylases.** Sarm1+/+ DRG neurons (top panels) or Sarm1+/- neurons (bottom panels) were cultured in the presence of vehicle (left panels) or of prolyl hydroxylase inhibitor Roxadustat (right panels), added at a concentration of 3 µg/ml on DIV 1 and re-supplemented every 2-3 days. Axotomy was performed on DIV 11, and the neurite integrity index (NII) was computed and plotted against time.

### Supplementary Fig. S4

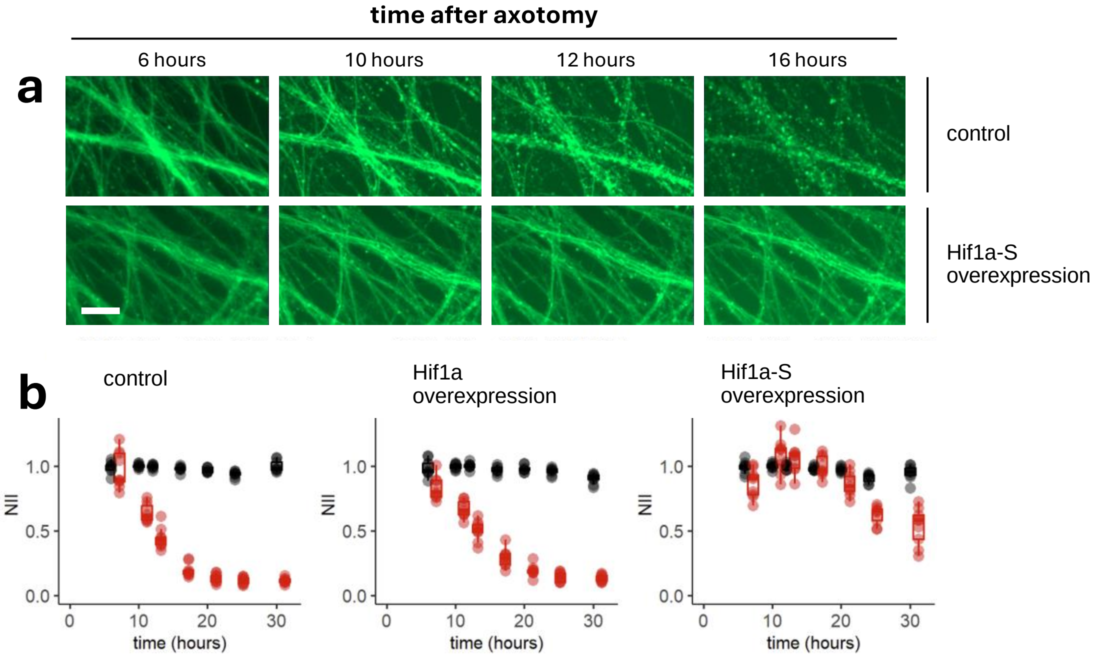


**Supplementary Fig. S4 | HIF-1 antagonizes Sarm1-mediated neurodegeneration**

**a,b** Axotomy experiment analogous to the experiment shown in Fig. 2a and b, with the difference that a slightly weaker expressing vector backbone was used. As a consequence, wild-type Hif1a is not protective (unlike in Fig. 2b), presumably because wild-type Hif1a protein ubiquitination and degradation is complete. Stabilized Hif1a-S robustly protects from axotomy-induced neurodegeneration.

### Supplementary Fig. S5

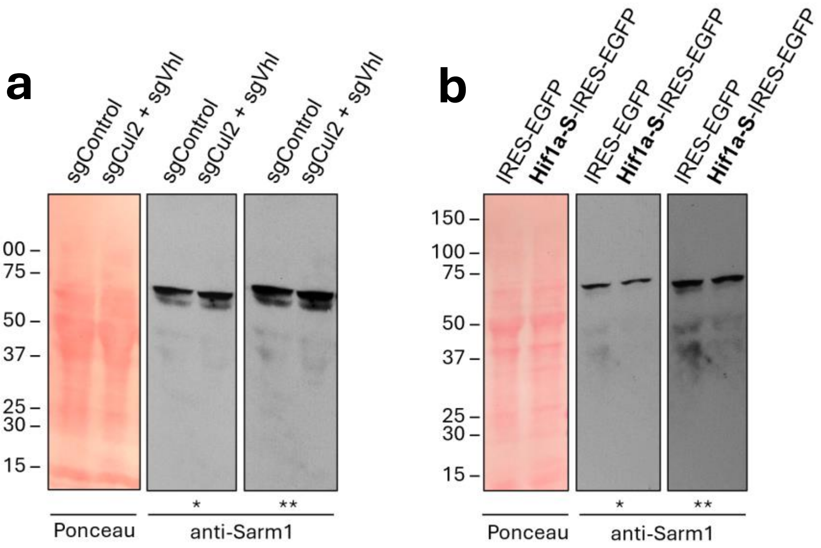


**Supplementary Fig. S5 | HIF-1 minimally alters Sarm1 protein levels in primary neurons**
**a** Mouse cortical neurons were transduced on DIV 1 with indicated lentiviral constructs. Cells were lysed on DIV 11, and proteins were electrophoretically separated and transferred onto nitrocellulose. Bulk protein was visualized using Ponceau S, and Sarm1 protein was detected by Western blot using chemiluminescence (* short exposure, ** long exposure).
**b** Western blot performed as in (a) using cells transduced with control vector (IRES-EGFP) or vector expressing Hif1a-S (Hif1a-S-IRES-EGFP).

### Supplementary Fig. S6

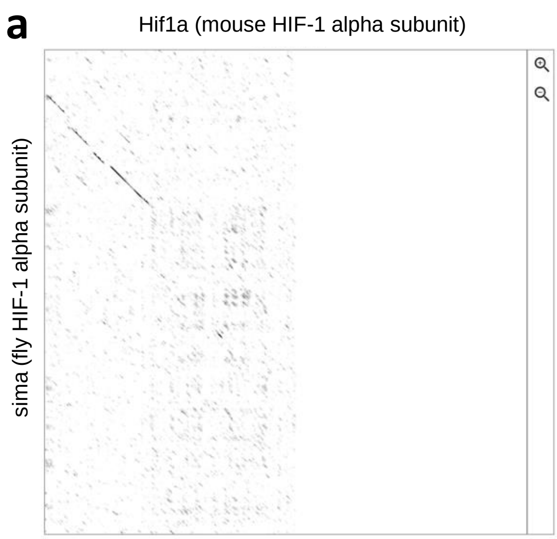


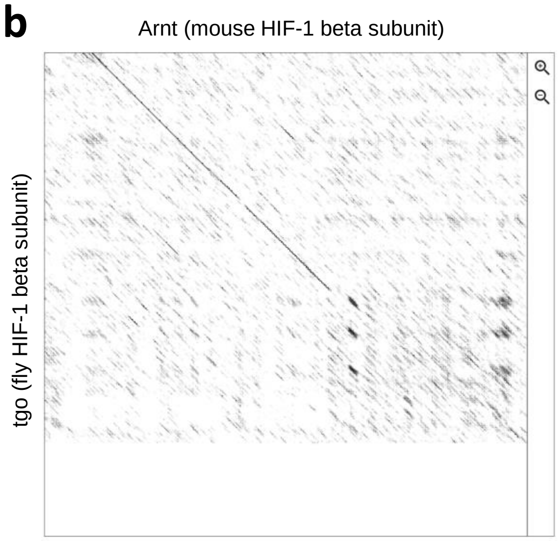


**Supplementary Fig. S6 | Protein sequence homologies between HIF-1 alpha and beta subunits from mouse and fly**

**a,b** Homology matrices were generated using Dotlet (https://dotlet.vital-it.ch/). Amino acid counts for compared proteins start in the upper left corner.

**a** Comparison of HIF-1 alpha chains from mouse (Hif1a) and fly (Sima)

**b** Comparison of HIF-1 beta chains from mouse (Arnt) and fly (Tgo, also known as tango)

### Supplementary Fig. S7

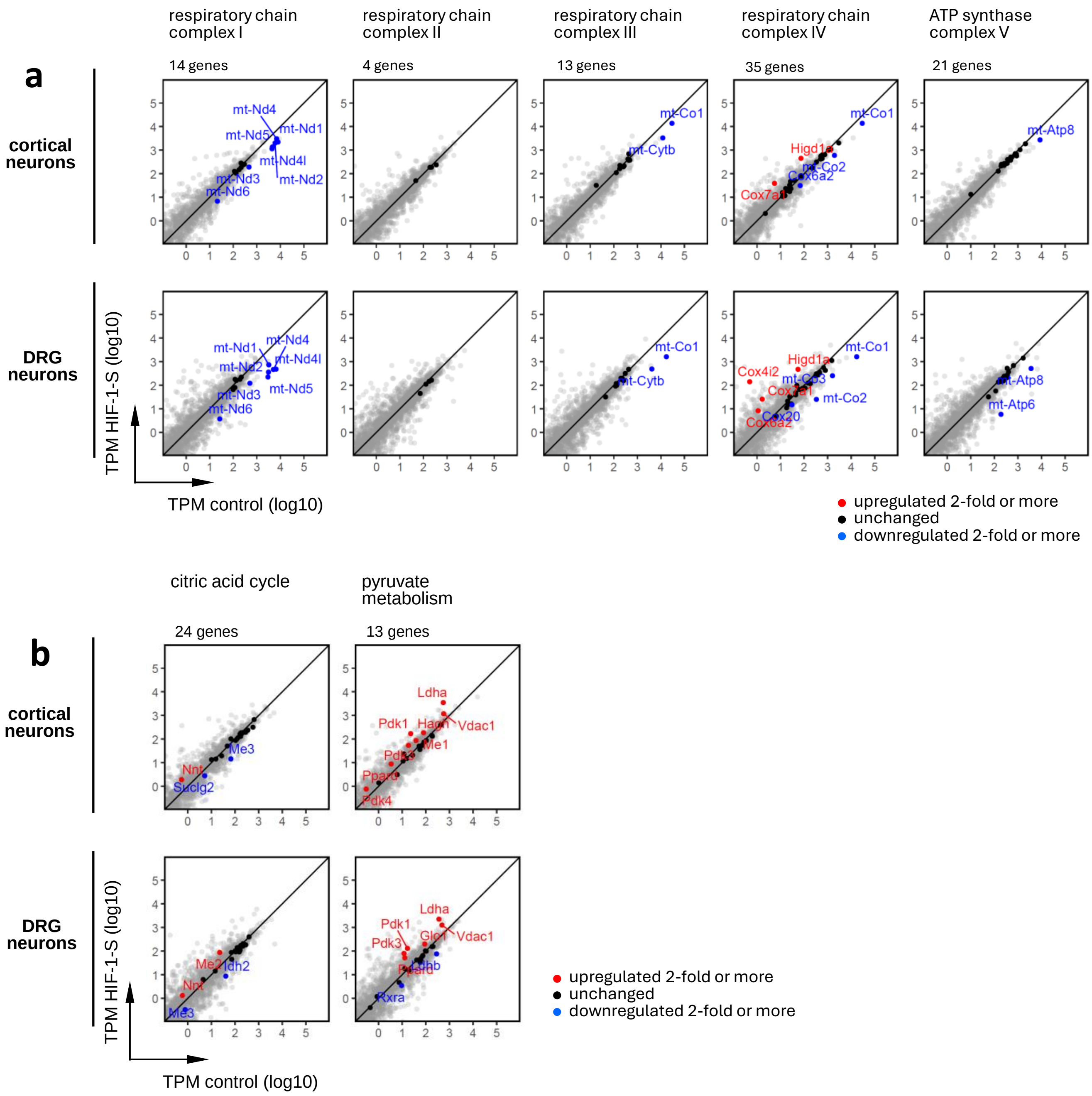


**Supplementary Fig. S7 | Transcriptional impact of HIF-1-S on major energy pathways in neurons**

**a,b** Red, genes upregulated 2-fold or more; blue, genes downregulated 2-fold or more; black, remaining genes. Gene numbers forming each complex or pathway are indicated. For comprehensive gene set information see Supplementary File “gene_sets”, tab “energy”.

**a** HIF-1-S-induced changes in expression of genes encoding the components of mitochondrial respiratory complexes I-IV, and of mitochondrial ATP synthase (complex V).

**b** HIF-1-S-mediated changes in gene expression for citric acid cycle and pyruvate metabolism.

### Supplementary Fig. S8

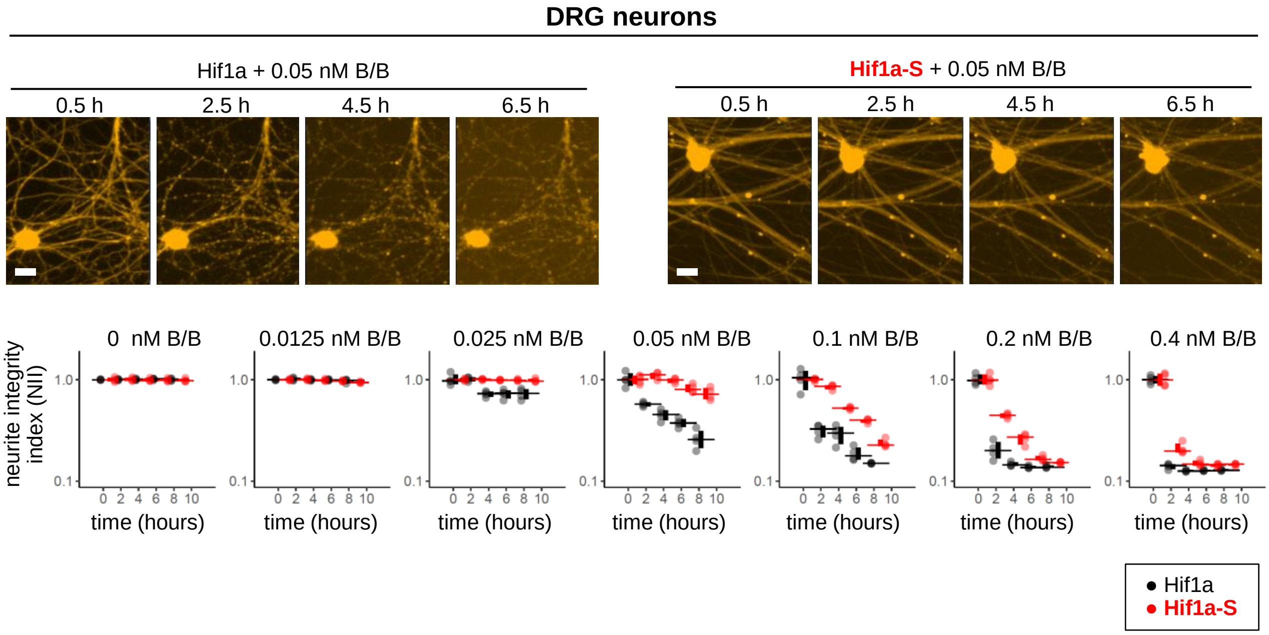


**Supplementary Fig. S8 | HIF-1 antagonizes neurodegeneration induced by Fh-TIR, a dimerization-activatable Sarm1 TIR domain**

Sparsely plated mouse DRG neurons were transduced on DIV 1 with wild-type Hif1a, or with degradation-resistant Hif1a-S. Fh-TIR (fused to mCherry) was included for all conditions. This experiment is analogous to the experiment in Fig. 7b, but compares wild-type Hif1a to Hif1a-S only, and has a finer grid of time points and B/B concentrations. For Fh-TIR glycohydrolase activation, B/B was added on DIV 11, and the neurite integrity index (NII) was computed and plotted against time (bottom panels).

### Supplementary Tables

#### Graphical data representation

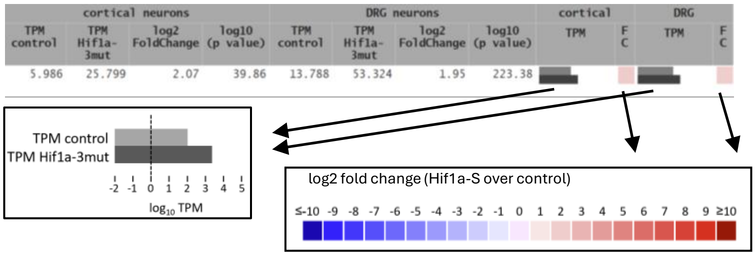


### T1 NAD+ biosynthesis

**sorted by**: cortical neurons, log2 fold change, high to low

| | | | | cortical neurons | | | | DRG neurons | | | | cortical | | DRG | |
|---|---|---|---|---|---|---|---|---|---|---|---|---|---|---|---|
| | gene symbol | gene description | gene type | TPM control | TPM Hif1a-3mut | log2 FoldChange | log10 (p value) | TPM control | TPM Hif1a-3mut | log2 FoldChange | log10 (p value) | TPM | F C | TPM | F C |
| 1 | Ido1 | indoleamine 2,3-dioxygenase 1 [Source:MGI Symbol;Acc:MGI:96416] | protein_coding | 0.193 | 1.286 | 2.77 | 1.45 | 0.216 | 0.158 | -0.45 | 0.12 | | | | |
| 2 | Ido2 | indoleamine 2,3-dioxygenase 2 [Source:MGI Symbol;Acc:MGI:2142489] | protein_coding | 0.816 | 4.638 | 2.47 | 6.10 | 0.220 | 3.312 | 3.89 | 14.40 | | | | |
| 3 | Nmrk2 | nicotinamide riboside kinase 2 [Source:MGI Symbol;Acc:MGI:1916814] | protein_coding | 0.184 | 0.664 | 1.87 | 0.58 | 0.199 | 0.249 | 0.29 | 0.09 | | | | |
| 4 | Qprt | quinolinate phosphoribosyltransferase [Source:MGI Symbol;Acc:MGI:1914625] | protein_coding | 0.026 | 0.114 | 1.77 | 0.20 | 0.126 | 0.131 | 0.08 | 0.02 | | | | |
| 5 | Haao | 3-hydroxyanthranilate 3,4-dioxygenase [Source:MGI Symbol;Acc:MGI:1349444] | protein_coding | 0.089 | 0.213 | 1.17 | 0.11 | 0.069 | 0.023 | -0.29 | 0.03 | | | | |
| 6 | Nadsyn1 | NAD synthetase 1 [Source:MGI Symbol;Acc:MGI:1926164] | protein_coding | 6.141 | 8.033 | 0.35 | 0.46 | 16.747 | 13.287 | -0.34 | 1.38 | | | | |
| 7 | Nmrk1 | nicotinamide riboside kinase 1 [Source:MGI Symbol;Acc:MGI:2147434] | protein_coding | 2.687 | 2.206 | -0.32 | 0.20 | 21.981 | 35.089 | 0.67 | 6.54 | | | | |
| 8 | Nmnat1 | nicotinamide nucleotide adenylyltransferase 1 [Source:MGI Symbol;Acc:MGI:1913704] | protein_coding | 4.051 | 3.121 | -0.39 | 0.36 | 9.130 | 11.055 | 0.27 | 0.82 | | | | |
| 9 | Tdo2 | tryptophan 2,3-dioxygenase [Source:MGI Symbol;Acc:MGI:1928486] | protein_coding | 0.895 | 0.717 | -0.40 | 0.11 | 0.078 | 0.000 | -0.85 | 0.08 | | | | |
| 10 | Nmnat2 | nicotinamide nucleotide adenylyltransferase 2 [Source:MGI Symbol;Acc:MGI:2444155] | protein_coding | 52.157 | 32.907 | -0.69 | 6.60 | 60.244 | 43.387 | -0.48 | 10.64 | | | | |
| 11 | Acmsd | amino carboxymuconate semialdehyde decarboxylase [Source:MGI Symbol;Acc:MGI:2386323] | protein_coding | 0.024 | 0.000 | -0.83 | 0.08 | 0.017 | 0.018 | 0.11 | 0.01 | | | | |
| 12 | Nmnat3 | nicotinamide nucleotide adenylyltransferase 3 [Source:MGI Symbol;Acc:MGI:1921330] | protein_coding | 1.818 | 0.868 | -0.94 | 0.51 | 0.399 | 1.377 | 1.80 | 1.97 | | | | |
| 13 | Afmid | arylformamidase [Source:MGI Symbol;Acc:MGI:2448704] | protein_coding | 4.717 | 1.805 | -1.39 | 1.98 | 5.578 | 12.907 | 1.21 | 6.13 | | | | |
| 14 | Kynu | kynureninase [Source:MGI Symbol;Acc:MGI:1918039] | protein_coding | 0.387 | 0.000 | -4.76 | 1.61 | 2.112 | 0.189 | -3.49 | 6.82 | | | | |
| 15 | Kmo | kynurenine 3-monooxygenase (kynurenine 3-hydroxylase) [Source:MGI Symbol;Acc:MGI:2138151] | protein_coding | 0.000 | 0.000 | | | 0.065 | 0.000 | -2.82 | 0.42 | | | | |

### T2 PARPs and Sirtuins

**sorted by**: gene symbol

| | | | | cortical neurons | | | | DRG neurons | | | | cortical | | DRG | |
|---|---|---|---|---|---|---|---|---|---|---|---|---|---|---|---|
| | gene symbol | gene description | gene type | TPM control | TPM Hif1a-3mut | log2 FoldChange | log10 (p value) | TPM control | TPM Hif1a-3mut | log2 FoldChange | log10 (p value) | TPM | F C | TPM | F C |
| 1 | Parp1 | poly (ADP-ribose) polymerase family, member 1 [Source:MGI Symbol;Acc:MGI:1340806] | protein_coding | 69.820 | 48.295 | -0.56 | 4.94 | 70.434 | 70.734 | 0.00 | 0.02 | | | | |
| 2 | Parp2 | poly (ADP-ribose) polymerase family, member 2 [Source:MGI Symbol;Acc:MGI:1341112] | protein_coding | 27.763 | 30.041 | 0.08 | 0.15 | 39.728 | 51.733 | 0.38 | 2.65 | | | | |
| 3 | Parp3 | poly (ADP-ribose) polymerase family, member 3 [Source:MGI Symbol;Acc:MGI:1891258] | protein_coding | 0.381 | 1.201 | 1.59 | 0.92 | 4.836 | 6.040 | 0.33 | 0.61 | | | | |
| 4 | Parp4 | poly (ADP-ribose) polymerase family, member 4 [Source:MGI Symbol;Acc:MGI:2685589] | protein_coding | 0.239 | 1.315 | 2.37 | 1.40 | 19.084 | 22.199 | 0.21 | 1.26 | | | | |
| 5 | Parp6 | poly (ADP-ribose) polymerase family, member 6 [Source:MGI Symbol;Acc:MGI:1914537] | protein_coding | 107.849 | 150.366 | 0.45 | 3.60 | 132.328 | 131.918 | -0.01 | 0.05 | | | | |
| 6 | Parp8 | poly (ADP-ribose) polymerase family, member 8 [Source:MGI Symbol;Acc:MGI:1098713] | protein_coding | 18.325 | 9.520 | -0.96 | 5.36 | 50.348 | 73.321 | 0.54 | 13.12 | | | | |
| 7 | Parp9 | poly (ADP-ribose) polymerase family, member 9 [Source:MGI Symbol;Acc:MGI:1933117] | protein_coding | 0.370 | 2.321 | 2.74 | 2.95 | 5.447 | 6.271 | 0.20 | 0.45 | | | | |
| 8 | Parp10 | poly (ADP-ribose) polymerase family, member 10 [Source:MGI Symbol;Acc:MGI:3712326] | protein_coding | 2.053 | 1.513 | -0.48 | 0.46 | 8.173 | 16.087 | 0.97 | 10.19 | | | | |
| 9 | Parp11 | poly (ADP-ribose) polymerase family, member 11 [Source:MGI Symbol;Acc:MGI:2141505] | protein_coding | 22.374 | 14.595 | -0.67 | 2.27 | 28.868 | 17.335 | -0.74 | 8.49 | | | | |
| 10 | Parp12 | poly (ADP-ribose) polymerase family, member 12 [Source:MGI Symbol;Acc:MGI:2143990] | protein_coding | 4.860 | 0.663 | -2.97 | 6.10 | 31.506 | 10.523 | -1.59 | 31.09 | | | | |
| 11 | Parp14 | poly (ADP-ribose) polymerase family, member 14 [Source:MGI Symbol;Acc:MGI:1919489] | protein_coding | 0.092 | 0.020 | -2.37 | 0.60 | 3.204 | 1.285 | -1.33 | 7.48 | | | | |
| 12 | Parp16 | poly (ADP-ribose) polymerase family, member 16 [Source:MGI Symbol;Acc:MGI:2446133] | protein_coding | 3.946 | 3.644 | -0.14 | 0.14 | 5.583 | 12.196 | 1.13 | 7.80 | | | | |
| 13 | Parpbp | PARP1 binding protein [Source:MGI Symbol;Acc:MGI:1922567] | protein_coding | 1.351 | 0.353 | -1.91 | 1.82 | 1.205 | 0.667 | -0.85 | 1.32 | | | | |
| 14 | Tiparp | TCDD-inducible poly(ADP-ribose) polymerase [Source:MGI Symbol;Acc:MGI:2159210] | protein_coding | 10.318 | 70.597 | 2.77 | 11.08 | 12.134 | 75.359 | 2.63 | 225.50 | | | | |
| 15 | Sirt1 | sirtuin 1 [Source:MGI Symbol;Acc:MGI:2135607] | protein_coding | 17.333 | 15.000 | -0.23 | 0.53 | 9.876 | 9.822 | -0.01 | 0.04 | | | | |
| 16 | Sirt2 | sirtuin 2 [Source:MGI Symbol;Acc:MGI:1927664] | protein_coding | 39.725 | 48.330 | 0.26 | 0.82 | 99.353 | 76.496 | -0.38 | 6.17 | | | | |
| 17 | Sirt3 | sirtuin 3 [Source:MGI Symbol;Acc:MGI:1927665] | protein_coding | 45.425 | 51.866 | 0.16 | 0.45 | 34.818 | 25.207 | -0.47 | 2.58 | | | | |
| 18 | Sirt4 | sirtuin 4 [Source:MGI Symbol;Acc:MGI:1922637] | protein_coding | 18.282 | 23.769 | 0.34 | 0.51 | 45.314 | 84.646 | 0.90 | 9.96 | | | | |
| 19 | Sirt5 | sirtuin 5 [Source:MGI Symbol;Acc:MGI:1915596] | protein_coding | 17.636 | 17.736 | -0.03 | 0.04 | 32.477 | 10.089 | -1.69 | 19.13 | | | | |
| 20 | Sirt6 | sirtuin 6 [Source:MGI Symbol;Acc:MGI:1354161] | protein_coding | 23.851 | 23.701 | -0.01 | 0.02 | 65.389 | 94.525 | 0.53 | 6.48 | | | | |
| 21 | Sirt7 | sirtuin 7 [Source:MGI Symbol;Acc:MGI:2385849] | protein_coding | 46.733 | 43.983 | -0.11 | 0.25 | 102.298 | 125.673 | 0.29 | 4.16 | | | | |

### T3 Complex IV subunit switch

**sorted by**: gene symbol

| | | | | cortical neurons | | | | DRG neurons | | | | cortical | | DRG | |
|---|---|---|---|---|---|---|---|---|---|---|---|---|---|---|---|
| | gene symbol | gene description | gene type | TPM control | TPM Hif1a-3mut | log2 FoldChange | log10 (p value) | TPM control | TPM Hif1a-3mut | log2 FoldChange | log10 (p value) | TPM | F C | TPM | F C |
| 1 | Cox4i1 | cytochrome c oxidase subunit 4I1 [Source:MGI Symbol;Acc:MGI:88473] | protein_coding | 1367.524 | 1080.374 | -0.36 | 2.04 | 669.247 | 557.530 | -0.27 | 4.21 | | | | |
| 2 | Cox4i2 | cytochrome c oxidase subunit 4I2 [Source:MGI Symbol;Acc:MGI:2135755] | protein_coding | 0.000 | 155.767 | 11.44 | 20.98 | 0.491 | 140.426 | 8.11 | 52.87 | | | | |
| 3 | Lonp1 | lon peptidase 1, mitochondrial [Source:MGI Symbol;Acc:MGI:1921392] | protein_coding | 47.944 | 122.534 | 1.33 | 22.12 | 76.647 | 343.550 | 2.16 | 321.00 | | | | |
| 4 | Lonp2 | lon peptidase 2, peroxisomal [Source:MGI Symbol;Acc:MGI:1914137] | protein_coding | 55.675 | 68.598 | 0.27 | 1.34 | 82.409 | 89.037 | 0.11 | 0.89 | | | | |

### Supplementary Tables (continued)

#### T4 Pentose phosphate pathway (MSigDB: REACTOME_PENTOSE_PHOSPHATE_PATHWAY)

**sorted by**: cortical neurons, log2 fold change, high to low

| | | | | cortical neurons | | | | DRG neurons | | | | cortical | | DRG | |
|---|---|---|---|---|---|---|---|---|---|---|---|---|---|---|---|
| | gene symbol | gene description | gene type | TPM control | TPM Hif1a-3mut | log2 FoldChange | log10 (p value) | TPM control | TPM Hif1a-3mut | log2 FoldChange | log10 (p value) | TPM | FC | TPM | FC |
| 1 | Dera | deoxyribose-phosphate aldolase (putative) [Source:MGI Symbol;Acc:MGI:1913762] | protein_coding | 0.823 | 2.761 | 1.75 | 1.86 | 4.959 | 8.226 | 0.75 | 1.88 | | | | |
| 2 | G6pdx | glucose-6-phosphate dehydrogenase X-linked [Source:MGI Symbol;Acc:MGI:105979] | protein_coding | 19.520 | 33.883 | 0.78 | 4.80 | 38.729 | 64.943 | 0.74 | 17.18 | | | | |
| 3 | Shpk | sedoheptulokinase [Source:MGI Symbol;Acc:MGI:1921887] | protein_coding | 2.178 | 3.392 | 0.61 | 0.69 | 7.326 | 7.544 | 0.04 | 0.07 | | | | |
| 4 | Tkt | transketolase [Source:MGI Symbol;Acc:MGI:105992] | protein_coding | 349.813 | 377.226 | 0.08 | 0.33 | 299.813 | 391.364 | 0.38 | 11.16 | | | | |
| 5 | Pgm2 | phosphoglucomutase 2 [Source:MGI Symbol;Acc:MGI:97564] | protein_coding | 11.618 | 12.222 | 0.04 | 0.06 | 15.202 | 34.332 | 1.17 | 18.59 | | | | |
| 6 | Pgd | phosphogluconate dehydrogenase [Source:MGI Symbol;Acc:MGI:97553] | protein_coding | 88.038 | 86.886 | -0.04 | 0.12 | 154.444 | 213.353 | 0.46 | 13.27 | | | | |
| 7 | Rpe | ribulose-5-phosphate-3-epimerase [Source:MGI Symbol;Acc:MGI:1913896] | protein_coding | 8.340 | 8.286 | -0.10 | 0.11 | 21.754 | 16.120 | -0.44 | 3.04 | | | | |
| 8 | Prps2 | phosphoribosyl pyrophosphate synthetase 2 [Source:MGI Symbol;Acc:MGI:97776] | protein_coding | 5.650 | 5.230 | -0.14 | 0.21 | 10.809 | 10.159 | -0.09 | 0.32 | | | | |
| 9 | Taldo1 | transaldolase 1 [Source:MGI Symbol;Acc:MGI:1274789] | protein_coding | 147.712 | 127.102 | -0.25 | 0.86 | 177.149 | 118.343 | -0.58 | 11.75 | | | | |
| 10 | Pgls | 6-phosphogluconolactonase [Source:MGI Symbol;Acc:MGI:1913421] | protein_coding | 48.763 | 39.600 | -0.33 | 0.89 | 92.564 | 69.972 | -0.41 | 5.03 | | | | |
| 11 | Prps1 | phosphoribosyl pyrophosphate synthetase 1 [Source:MGI Symbol;Acc:MGI:97775] | protein_coding | 60.935 | 41.481 | -0.59 | 3.56 | 75.790 | 37.700 | -1.01 | 26.85 | | | | |
| 12 | Rpia | ribose 5-phosphate isomerase A [Source:MGI Symbol;Acc:MGI:103254] | protein_coding | 19.844 | 11.714 | -0.79 | 2.67 | 15.897 | 16.590 | 0.06 | 0.14 | | | | |
| 13 | Rbks | ribokinase [Source:MGI Symbol;Acc:MGI:1918586] | protein_coding | 2.044 | 0.639 | -1.78 | 0.80 | 2.021 | 1.370 | -0.56 | 0.39 | | | | |

#### T5 Glycolytic enzymes, glucose transporters, and lactate dehydrogenase

**sorted by**: cortical neurons, log2 fold change, high to low

| | | | | cortical neurons | | | | DRG neurons | | | | cortical | | DRG | |
|---|---|---|---|---|---|---|---|---|---|---|---|---|---|---|---|
| | gene symbol | gene description | gene type | TPM control | TPM Hif1a-3mut | log2 FoldChange | log10 (p value) | TPM control | TPM Hif1a-3mut | log2 FoldChange | log10 (p value) | TPM | FC | TPM | FC |
| 1 | Hk2 | hexokinase 2 [Source:MGI Symbol;Acc:MGI:1315197] | protein_coding | 2.307 | 435.209 | 7.48 | 15.64 | 3.254 | 390.461 | 6.91 | 153.05 | | | | |
| 2 | Slc2a1 | solute carrier family 2 (facilitated glucose transporter), member 1 [Source:MGI Symbol;Acc:MGI:95755] | protein_coding | 34.335 | 1289.655 | 5.21 | 268.92 | 51.830 | 1427.672 | 4.78 | 345.00 | | | | |
| 3 | Aldoc | aldolase C, fructose-bisphosphate [Source:MGI Symbol;Acc:MGI:101863] | protein_coding | 19.961 | 604.813 | 4.89 | 339.00 | 92.014 | 508.335 | 2.46 | 198.32 | | | | |
| 4 | Pfkfb3 | 6-phosphofructo-2-kinase/fructose-2,6-biphosphatase 3 [Source:MGI Symbol;Acc:MGI:2181202] | protein_coding | 19.540 | 178.634 | 3.18 | 106.71 | 36.176 | 315.757 | 3.12 | 338.00 | | | | |
| 5 | Pfkl | phosphofructokinase, liver, B-type [Source:MGI Symbol;Acc:MGI:97547] | protein_coding | 46.269 | 354.619 | 2.91 | 158.61 | 109.702 | 545.353 | 2.31 | 344.00 | | | | |
| 6 | Gpi1 | glucose-6-phosphate isomerase 1 [Source:MGI Symbol;Acc:MGI:95797] | protein_coding | 599.859 | 3954.901 | 2.70 | 161.45 | 1321.700 | 5027.140 | 1.92 | 327.00 | | | | |
| 7 | Ldha | lactate dehydrogenase A [Source:MGI Symbol;Acc:MGI:96759] | protein_coding | 551.277 | 3496.562 | 2.64 | 121.49 | 361.524 | 2236.152 | 2.63 | 349.00 | | | | |
| 8 | Slc2a3 | solute carrier family 2 (facilitated glucose transporter), member 3 [Source:MGI Symbol;Acc:MGI:95757] | protein_coding | 358.021 | 2158.446 | 2.57 | 125.24 | 195.364 | 2249.789 | 3.52 | 339.00 | | | | |
| 9 | Pgk1 | phosphoglycerate kinase 1 [Source:MGI Symbol;Acc:MGI:97555] | protein_coding | 456.523 | 2525.775 | 2.44 | 93.31 | 301.548 | 977.541 | 1.69 | 137.13 | | | | |
| 10 | Tpi1 | triosephosphate isomerase 1 [Source:MGI Symbol;Acc:MGI:98797] | protein_coding | 333.600 | 1701.684 | 2.32 | 66.68 | 367.060 | 1283.996 | 1.80 | 193.10 | | | | |
| 11 | Gapdh | glyceraldehyde-3-phosphate dehydrogenase [Source:MGI Symbol;Acc:MGI:95640] | protein_coding | 2113.993 | 9857.280 | 2.20 | 69.85 | 2220.847 | 9057.448 | 2.02 | 173.79 | | | | |
| 12 | Hk3 | hexokinase 3 [Source:MGI Symbol;Acc:MGI:2670962] | protein_coding | 0.022 | 0.093 | 2.12 | 0.30 | 0.000 | 0.039 | 2.53 | 0.31 | | | | |
| 13 | Aldoa | aldolase A, fructose-bisphosphate [Source:MGI Symbol;Acc:MGI:87994] | protein_coding | 936.994 | 3164.815 | 1.73 | 53.20 | 1457.092 | 3363.286 | 1.20 | 134.09 | | | | |
| 14 | Aldob | aldolase B, fructose-bisphosphate [Source:MGI Symbol;Acc:MGI:87995] | protein_coding | 1.098 | 3.467 | 1.65 | 1.77 | 1.205 | 1.412 | 0.23 | 0.17 | | | | |
| 15 | Pkm | pyruvate kinase, muscle [Source:MGI Symbol;Acc:MGI:97591] | protein_coding | 821.701 | 2461.299 | 1.55 | 45.87 | 653.352 | 3689.199 | 2.49 | 349.00 | | | | |
| 16 | Hk1 | hexokinase 1 [Source:MGI Symbol;Acc:MGI:96103] | protein_coding | 248.808 | 673.705 | 1.41 | 40.97 | 450.002 | 802.086 | 0.83 | 64.81 | | | | |
| 17 | Eno1 | enolase 1, alpha non-neuron [Source:MGI Symbol;Acc:MGI:95393] | protein_coding | 613.858 | 1438.044 | 1.20 | 23.29 | 934.902 | 3458.736 | 1.88 | 248.53 | | | | |
| 18 | Eno1b | enolase 1B, retrotransposed [Source:MGI Symbol;Acc:MGI:3648653] | protein_coding | 1.298 | 2.954 | 1.14 | 0.96 | 2.257 | 7.454 | 1.71 | 5.90 | | | | |
| 19 | Hkdc1 | hexokinase domain containing 1 [Source:MGI Symbol;Acc:MGI:2384910] | protein_coding | 0.930 | 2.047 | 1.10 | 1.40 | 0.217 | 0.275 | 0.84 | 0.30 | | | | |
| 20 | Aldoart1 | aldolase 1 A, retrogene 1 [Source:MGI Symbol;Acc:MGI:2447811] | protein_coding | 0.000 | 0.038 | 1.09 | 0.10 | 0.000 | 0.019 | 1.07 | 0.10 | | | | |
| 21 | Pfkm | phosphofructokinase, muscle [Source:MGI Symbol;Acc:MGI:97548] | protein_coding | 210.143 | 251.617 | 0.23 | 1.63 | 157.601 | 121.272 | -0.38 | 9.38 | | | | |
| 22 | Gck | glucokinase [Source:MGI Symbol;Acc:MGI:1270854] | protein_coding | 2.432 | 2.800 | 0.09 | 0.05 | 2.502 | 3.913 | 0.65 | 1.12 | | | | |
| 23 | Eno2 | enolase 2, gamma neuronal [Source:MGI Symbol;Acc:MGI:95394] | protein_coding | 632.926 | 674.753 | 0.06 | 0.25 | 414.476 | 920.893 | 1.15 | 125.11 | | | | |
| 24 | Eno3 | enolase 3, beta muscle [Source:MGI Symbol;Acc:MGI:95395] | protein_coding | 7.076 | 7.183 | -0.03 | 0.02 | 17.929 | 9.337 | -0.94 | 4.23 | | | | |
| 25 | Gapdhrt | glyceraldehyde-3-phosphate dehydrogenase, retrotransposed [Source:MGI Symbol;Acc:MGI:3782011] | protein_coding | 0.301 | 0.232 | -0.20 | 0.04 | 0.481 | 0.104 | -2.36 | 1.22 | | | | |
| 26 | Pfkfb2 | 6-phosphofructo-2-kinase/fructose-2,6-biphosphatase 2 [Source:MGI Symbol;Acc:MGI:107815] | protein_coding | 24.133 | 20.574 | -0.27 | 0.63 | 37.532 | 31.489 | -0.26 | 2.58 | | | | |
| 27 | Bpgm | 2,3-bisphosphoglycerate mutase [Source:MGI Symbol;Acc:MGI:1098242] | protein_coding | 28.314 | 23.545 | -0.31 | 0.76 | 23.193 | 8.228 | -1.50 | 23.48 | | | | |
| 28 | Pgam5 | phosphoglycerate mutase family member 5 [Source:MGI Symbol;Acc:MGI:1919792] | protein_coding | 66.875 | 52.999 | -0.36 | 1.85 | 47.686 | 40.617 | -0.23 | 1.71 | | | | |
| 29 | Aldoa | aldolase A, fructose-bisphosphate [Source:MGI Symbol;Acc:MGI:87994] | protein_coding | 7.497 | 5.405 | -0.46 | 0.29 | 1.472 | 10.143 | 2.77 | 7.50 | | | | |
| 30 | Eno4 | enolase 4 [Source:MGI Symbol;Acc:MGI:2441717] | protein_coding | 0.663 | 0.393 | -0.82 | 0.30 | 0.864 | 0.494 | -0.83 | 0.58 | | | | |
| 31 | Pfkfb4 | 6-phosphofructo-2-kinase/fructose-2,6-biphosphatase 4 [Source:MGI Symbol;Acc:MGI:2687284] | protein_coding | 26.372 | 13.821 | -0.97 | 5.77 | 16.477 | 12.075 | -0.45 | 2.36 | | | | |
| 32 | Pfkfb1 | 6-phosphofructo-2-kinase/fructose-2,6-biphosphatase 1 [Source:MGI Symbol;Acc:MGI:107816] | protein_coding | 4.483 | 2.233 | -1.02 | 1.01 | 4.770 | 3.137 | -0.60 | 1.01 | | | | |
